# BAF complexes maintain accessibility at stimulus-responsive chromatin and are required for transcriptional stimulus responses

**DOI:** 10.64898/2026.03.19.712964

**Authors:** Alexander O. D. Gulka, Kihoon A. Kang, Ziben Zhou, David U. Gorkin

## Abstract

**Background:** Gene expression changes in response to developmental and environmental cues rely on *cis*-regulatory sequence elements (cREs). BRG1/BRM-Associated Factors (BAF) chromatin remodeling complexes maintain chromatin accessibility at many cREs, enabling binding by transcription factors (TFs). However, cREs exhibit a broad range of sensitivity to loss of BAF function, and the basis of this variability remains unknown.

**Results:** To identify the characteristics of BAF-dependent cREs, we mapped chromatin accessibility changes following acute pharmacologic BAF inhibition in GM12878 lymphoblastoid cells. We integrated these results with over 100 TF and histone modification ChIP-seq datasets and used machine learning to identify features that predict chromatin accessibility changes. We found that Activator Protein 1 (AP-1) factors and lymphoid lineage-defining TFs including RUNX3 and PU.1 predicted BAF-dependence. Strikingly, we found that cREs bearing the chromatin signature of “primed” enhancers – enriched for H3K4me1 but lacking H3K27ac – were significantly more sensitive to BAF inhibition than typical active enhancers. As primed enhancers are known to facilitate transcriptional responses to stimuli, we tested the requirement of BAF activity in these responses. Acute BAF inhibition was sufficient to prevent both chromatin and transcriptional responses to interferon gamma and dexamethasone. cREs which normally gained accessibility in response to stimulation failed to do so with BAF inhibition, and these cREs were linked to genes with suppressed transcriptional induction.

**Conclusions:** Collectively, our results demonstrate a requirement for continuous BAF activity to enable stimulus response and suggest that defective signal responsiveness may be a pathogenic mechanism in disease states caused by loss-of-function mutations in BAF subunits.

## BACKGROUND

The cellular response to external cues involves molecular changes at *cis*-regulatory elements (cREs) [1–3]. Chromatin remodeling is an early and critical step in the activation of cREs, whereby local changes to DNA packaging generate an “accessible” chromatin state that is permissive to binding by transcription factors (TFs) and other regulatory proteins [4, 5]. In mammals, BRG1/BRM-Associated Factors (BAF; also known as mammalian SWI/SNF) complexes regulate chromatin accessibility at cREs. BAF complexes use energy from ATP hydrolysis to reposition nucleosomes, thus increasing DNA accessibility to binding factors [6]. Consistent with the central role of BAF complexes in transcriptional regulation, mutations in genes encoding BAF subunits are linked to a variety of human diseases. Mutations in any one of 15 BAF complex subunits can cause Mendelian developmental disorders, collectively referred to as SWI/SNF-related Intellectual Disability Disorders [7]. In addition, somatic mutations in BAF subunits are observed in a spectrum of cancers [8–10], and are hallmark driver events in a set of rare pediatric malignancies [9, 11]. Notably, many disease-causing BAF mutations are predicted to result in loss-of-function [12]. Thus, defining the molecular consequences of reduced BAF function has broad implications for understanding how BAF mutations cause disease.

The chromatin remodeling function of BAF complexes depends on ATP hydrolysis, mediated by one of two mutually exclusive paralogous ATPase subunits, SMARCA2 and SMARCA4. Small molecule approaches targeting these subunits have been developed including SMARCA2/A4 inhibitors [13] and Proteolysis-Targeting Chimeras (PROTACs) that induce degradation of these subunits [14]. Importantly, the changes to chromatin accessibility resulting from inhibition or PROTAC-induced degradation closely parallel those observed with genetically encoded degron systems (SMARCA4-dTAG, SMARCA4-AID) and loss-of-function genetic mutations in SMARCA2/4 [15–18]. Prior studies using these small molecules have established that inactivation of BAF complexes results in rapid loss of chromatin accessibility at thousands of cREs [15, 16]. However, the requirement for BAF to maintain chromatin accessibility (hereafter referred to as “BAF-dependence”) varies widely across cREs. Enhancers are the most BAF-dependent across a variety of cell types [15, 16, 18, 19], but, even among enhancers, there is striking variability in BAF-dependence. Enhancers can be subcategorized based on many factors, including their histone modification signatures, the TFs that bind them, the types of genes they activate, and the stimuli that they respond to. For example, cells maintain repertoires of both “active” enhancers (H3K4me1+/H3K27ac+)[20, 21], which are thought to direct ongoing gene expression, and “primed” enhancers, which can rapidly be activated in response to specific stimuli. Primed enhancers are characterized by the presence of accessible chromatin and H3K4me1, but they have only weak or absent H3K27ac and are thought to lack the full complement of TFs and coactivators necessary for regulatory activity [20, 22, 23].

In this study we sought to define the basis of variable BAF-dependence among enhancers. We made use of the lymphoblastoid cell line GM12878, in which ChIP-seq data is available to map the binding locations of more than 100 TFs and nearly a dozen histone modifications. We hypothesized that specific patterns of TF binding and histone modification could distinguish BAF-dependent and -independent enhancers. To measure the dependence of cREs on BAF activity, we used both ATPs inhibitors and PROTAC degraders of SMARCA2/4 and generated time- and dosage-response maps of chromatin accessibility with the assay for transposase-accessible chromatin followed by sequencing (ATAC-seq).We then applied machine learning to identify TFs and histone modifications predictive of chromatin accessibility changes after BAF inhibition. We found that multiple TFs are strongly predictive of BAF-dependence, including AP-1 factors, RUNX3, and PU.1. Moreover, we found that primed enhancers are distinctly sensitive to BAF inhibition. This finding led us to further investigate the role of BAF in priming transcriptional responses to external stimuli. Strikingly, we found that the chromatin and transcriptional responses to two orthogonal stimuli - interferon gamma and dexamethasone - require continuous BAF activity. Stimulus-responsive enhancers are exceptionally sensitive to BAF inhibition and physically interact with stimulus-responsive genes that fail to activate. Taken together, our results reveal specific enhancer features that dictate BAF-dependence and demonstrate that diverse stimulus responses rely on continuous BAF activity. These findings point to molecular mechanisms, including chromatin-based stimulus response, that may underlie the roles of BAF subunit mutations in a variety of disease states.

## RESULTS

### BAF inhibition causes widespread but variable loss of chromatin accessibility at enhancers

To identify BAF-dependent cREs we comprehensively profiled the effects of BAF inhibition on chromatin accessibility in the lymphoblastoid cell line GM12878 (**Figure 1A**). We selected BRM014, an allosteric inhibitor of SMARCA2 and SMARCA4, as our primary means of BAF inhibition due to its demonstrated potency and specificity. BRM014 has been shown to alter chromatin accessibility within minutes of treatment [15], and has similar effects to genetically-encoded degron systems (SMARCA4-dTAG, SMARCA4-AID) and proteolysis-targeting chimeras against SMARCA2/4 [15, 17, 18, 24]. Hereafter we will use the label “BAF^inh^” to refer to BAF inhibition using BRM014. We confirmed that no cellular viability defects were observed with BAF^inh^ doses up to 10µM and treatment durations up to 24 hours (**Supplemental Figure 1A**). This concentration is in line with doses used in published reports with this compound [15, 16, 18]. We first treated cells for 0.5, 1, 2, 6, and 24h with 10µM BAF^inh^ and measured chromatin accessibility by ATAC-seq. Strikingly, we observed widespread reduction in accessibility after only 0.5h of treatment, with 52,207 ATAC-seq peaks significantly losing accessibility (FDR < 0.05, log2FC < -1; **Figure 1B, C; Supplemental Table 1**). The number of peaks losing accessibility (i.e. “BAF-dependent” peaks) remained relatively stable over the time course, whereas gained peaks - though far fewer - grew with treatment duration. This suggests that the primary and immediate effect of BAF^inh^ is accessibility loss, while accessibility gains may be secondary, consistent with published reports [17]. Similar patterns were observed across a range of doses (10nM, 100nM, 1µM, 10µM) at 6 hours of treatment (**Figure 1B, C**). BAF^inh^ effects on chromatin accessibility were overall highly concordant between doses and timepoints (**Supplemental Figure 1B**). To confirm the specificity of the effects on accessibility observed with BRM014, we treated cells with ACBI1, a proteolysis-targeting chimera targeting SMARCA2 and SMARCA4 [14]. While the effect sizes with this approach were more modest, likely due to incomplete clearance of SMARCA2 (**Supplemental Figure 1C**) we observed high concordance between cREs affected by ACBI1 and BRM014 **(Supplemental Figure 1D, E**). Notably, while our results demonstrate that accessible chromatin in GM12878 is broadly dependent on BAF activity, many regions do not show significant change in accessibility even after 24 hours of 10µM BRM014 treatment (n = 53,917).

**Figure 1:**
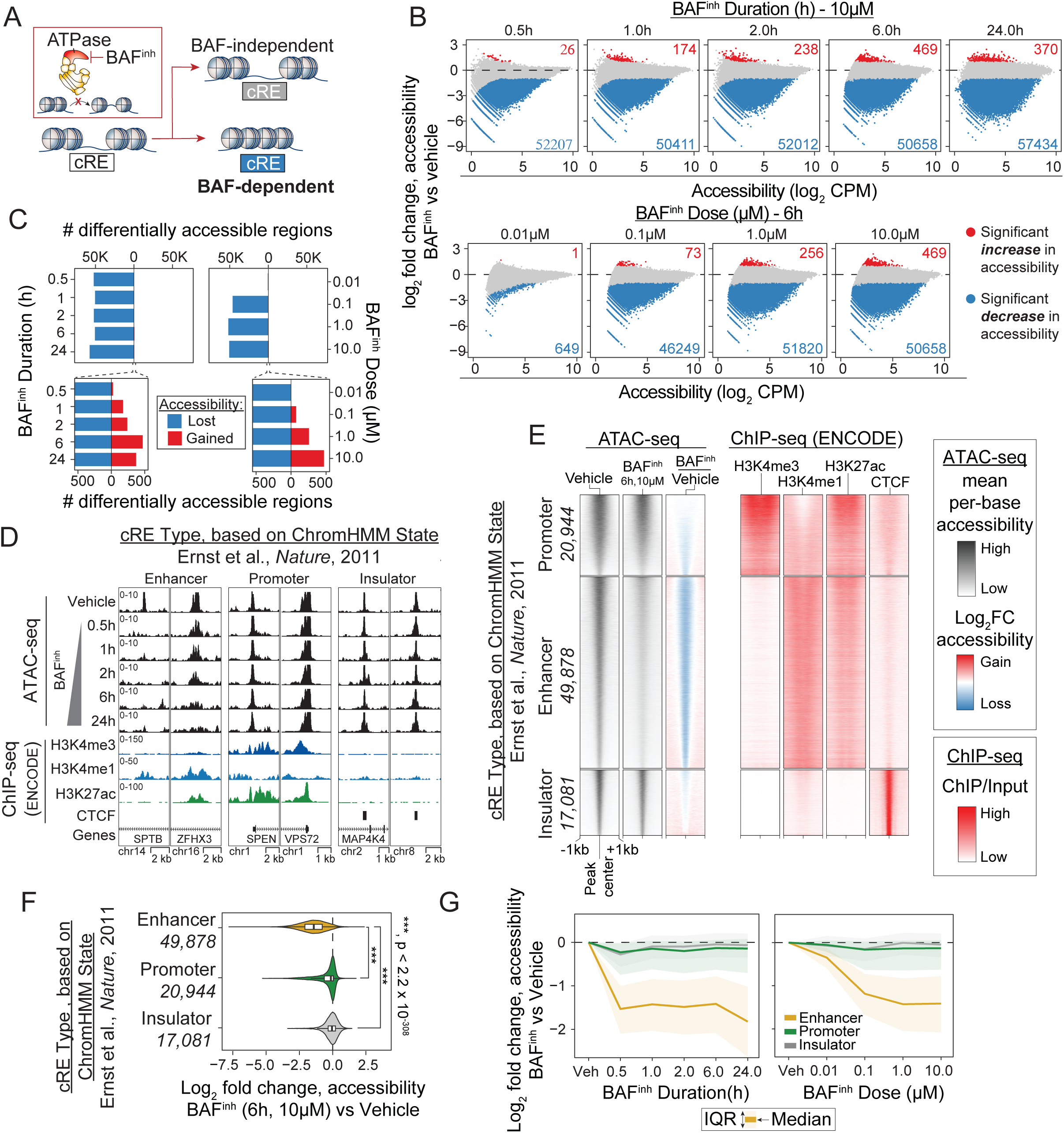
BAF^inh^ leads to widespread reduction in cRE accessibility, especially at enhancers. (A) Schematic of approach. The allosteric SMARCA2/A4 inhibitor BRM014 (“BAF^inh^”) is used to define BAF-dependent *cis*-regulatory elements (cREs) at different timepoints and dosages. (B) Accessibility change versus mean accessibility (MA) plots showing effects of BAF^inh^ on chromatin accessibility across all GM12878 ATAC-seq peaks (n = 111,721). Numbers of peaks with significantly increased (FDR < .05, log2 fold change > 1) or decreased (FDR < .05, log2 fold change < -1 accessibility are indicated. (C) Chromatin accessibility is predominantly lost upon BAF^inh^, with only gradual accumulation accessibility gains at later times and higher doses. (D) Representative genome browser examples of enhancers, promoters, and insulators illustrating greater BAF-dependence of enhancers. cRE annotations were derived from a published 15-state ChromHMM model [25]. (E) Enhancers, but not promoters or insulators, exhibit widespread loss of accessibility following BAF^inh^ (6h, 10µM). (F) Enhancers lose significantly more accessibility following BAF^inh^ (6h, 10µM) than promoters or insulators. P-values for each comparison were calculated using a Wilcoxon rank-sum test. (G) Enhancers lose accessibility more rapidly (left) and at lower BAF^inh^ doses (right) than other cRE classes.

Dependence on BAF activity varies between classes of cREs [15, 16, 18]. To evaluate whether similar patterns are seen in GM12878, we leveraged published chromatin state annotations to assign cRE types to ATAC-seq peaks (**Figure 1B, Supplemental Figure 1F)** [25]. In line with reports in other cell types, we found that GM12878 enhancers are more BAF-dependent than promoters or CTCF-bound (“insulator”) elements (**Figure 1D-F**). Enhancers are significantly enriched within BAF-dependent peaks (OR = 15.0, p < 2.2 x 10^-308^, Fisher’s exact test), while promoters and CTCF-bound elements are significantly depleted from BAF-dependent peaks (**Supplemental Figure 1G**). Strikingly, 71% and 65% of enhancers significantly lose accessibility by 0.5h (10µM) and 100nM (6h) inhibitor treatment, respectively, highlighting the role of BAF complexes in regulating enhancer accessibility (**Supplemental Figure 1H)**. We further evaluated the time- and dose-response to BAF^inh^ and found that enhancers lose accessibility more rapidly and at a lower dose of inhibitor than other cRE types (**Figure 1G**). To ensure that our findings were robust to cRE annotation approach, we reclassified cREs using two alternative methods: (1) ENCODE candidate cis-regulatory element (cCRE) annotations [26], and (2) using publicly available H3K4me1, H3K4me3, and CTCF ChIP-seq data to annotate cREs, taking advantage of well-established differences in these marks between cRE types [20, 21, 27]. We observed similar patterns of BAF-dependence with both alternative annotations (**Supplemental Figure 1I, J**). For ease of interpretation, we utilize the ChIP-based annotations (**Supplemental Figure 1J**) throughout the remainder of this study. Together, these results demonstrate the critical role of BAF in maintaining chromatin accessibility in GM12878, as in other cell types, and show that enhancers are especially BAF-dependent. We next sought to leverage the rich epigenomic data series unique to GM12878 to more deeply dissect features underlying enhancer BAF-dependence.

### Machine learning identifies chromatin features associated with BAF-dependence

Although enhancers display higher average BAF dependence than other cRE classes, we observed considerable variability in BAF-dependence among enhancers (**Figure 2A, B; Supplemental Figure 2A**). We reasoned that molecularly and/or functionally distinct enhancer subsets may be differentially reliant on BAF activity, such that chromatin features could predict BAF-dependence. To test this, we developed a machine learning pipeline that integrates random forest classification for prediction of binary chromatin accessibility outcomes with ridge regression to model quantitative accessibility changes based on ChIP-seq data (**Figure 2C, Supplemental Figure 2B)**.

**Figure 2:**
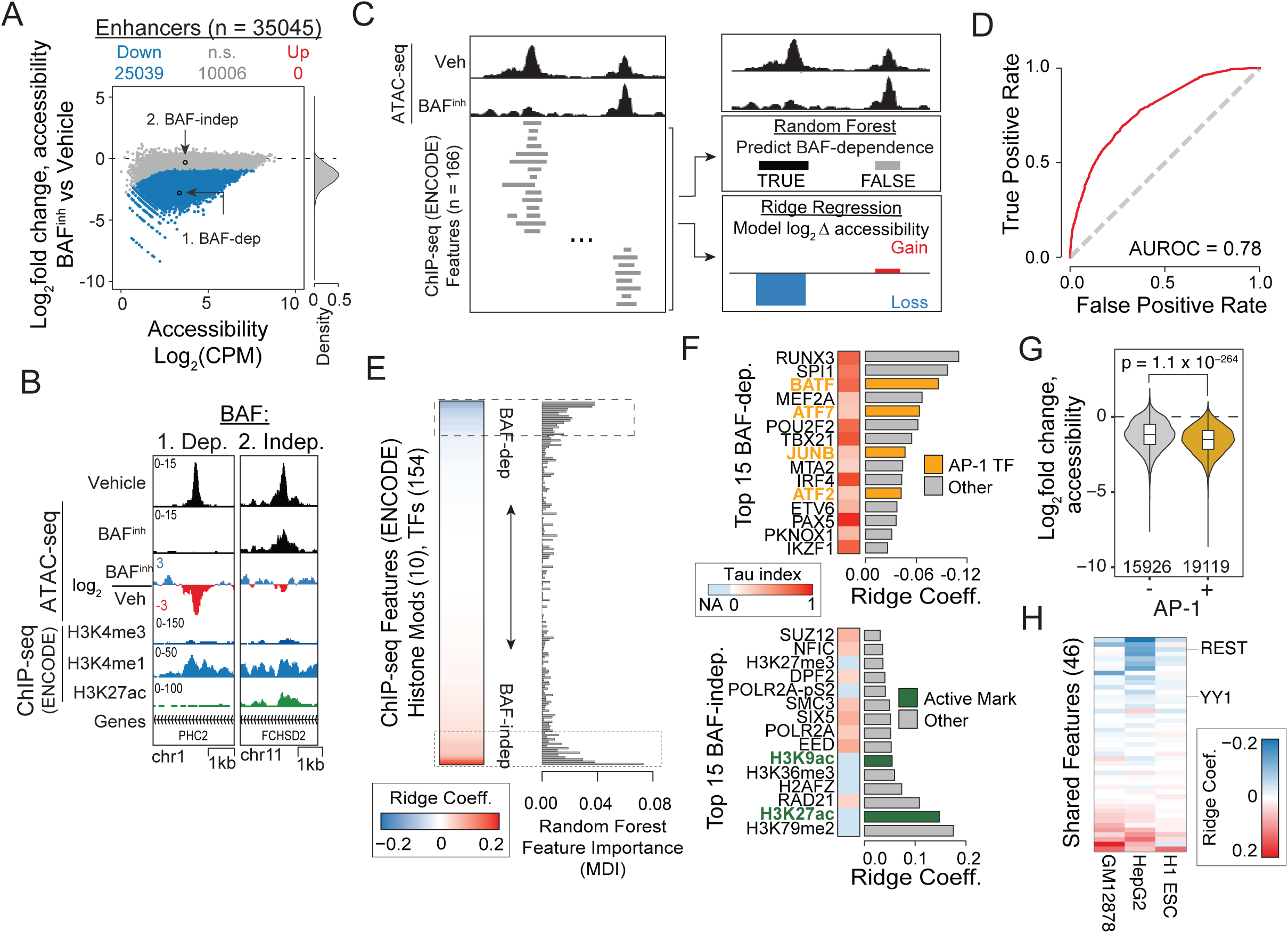
Machine learning identifies distinct chromatin features of BAF-dependent enhancers. (A) MA plot (left) and Log2 accessibility fold change density distribution (right) demonstrating variation in sensitivity to BAF^inh^ (0.5h, 10µM) among enhancers. Representative BAF-dependent and independent enhancers are indicated and shown in more detail in (B). (B) Genome browser examples of BAF-dependent and -independent enhancers from (A). (C) Conceptual representation of machine learning framework. ATAC-seq peaks were annotated based on overlap with ENCODE ChIP-seq peaks for 155 transcription factors and 11 histone modifications (left). This feature matrix was used to predict binary BAF dependence using a random forest classifier (top right) and to model continuous accessibility log2 fold changes using ridge-regularized regression (bottom right). (D) The random forest model performs strongly at predicting BAF-dependence among enhancers. (E) Machine learning identifies chromatin features predictive of both BAF-dependence and independence. The top 15 BAF-dependence and -independence features are indicated by dashed boxes. H3K4me3 and CTCF are excluded by design in enhancer-only models, resulting in a total of 154 transcription factor and 10 histone modification features. (F) Features associated with BAF dependence include multiple AP-1 transcription factors and show tissue-restricted expression patterns, while BAF-independence features include activating chromatin marks and features with more ubiquitous expression. Tissue-specificity was quantified using the tau (τ) index, where τ = 0 represents ubiquitous expression and τ = 1 corresponds to expression in a single tissue (see **Methods**). (G) AP-1 bound enhancers are significantly more BAF-dependent than non-bound enhancers. P-value calculated using a Wilcoxon rank-sum test. (H) Application of machine learning approach across multiple cell types (GM12878, HepG2, H1 embryonic stem cells) identifies shared features associated with BAF-dependence, including REST and YY1.

We leveraged 166 publicly available ChIP-seq datasets in GM12878 - comprising 11 histone PTMs and 155 TFs (**Supplemental Table 2**) - and asked whether we could predict accessibility changes due to BAF^inh^ based on these features. When applied to the total set of GM12878 cREs, the random forest classifier displayed strong predictive performance on a held-out test set (AUROC = 0.89; **Supplemental Figure 2C**). Moreover, feature importance in the random forest model, as measured by mean decrease in impurity (MDI), was highly correlated with ridge regression coefficients (r = 0.696, p = 2.5 x 10^-25^; **Supplemental Figure 2D**), indicating high consistency in feature contributions (i.e. which chromatin features were most informative) to both binary and quantitative models. Our approach is robust to choice of random forest feature importance metric, as we observed strong agreement between MDI and permutation-based feature importance scores (r = 0.92, p = 2.5 x 10^-68^; **Supplemental Figure 2E**). To further evaluate our approach, we trained independent models for all BAF^inh^ conditions (four doses and five timepoints) and compared their performance and feature metrics. Our method achieved consistent performance across conditions, and both feature importance metrics were highly correlated between models (**Supplemental Figure 2F**). In agreement with our earlier findings, models consistently identified CTCF and promoter features such as H3K4me3 and Pol II among the most predictive features of BAF-independence, while the enhancer-associated histone mark H3K4me1 was associated with BAF-dependence (**Supplemental Figure 2G**). Collectively, these results demonstrate the utility and robustness of our machine learning approach as a tool to identify chromatin features that predict perturbation outcomes.

We next sought to identify chromatin features that are highly predictive of BAF-dependence specifically among enhancers, excluding elements overlapping H3K4me3 or CTCF peaks, or within 1kb of an annotated TSS from our input feature matrix. We found that prediction performance was only moderately reduced with this limited input set (AUROC = 0.78, **Figure 2D**). Similar to the model trained on the full set of ATAC-seq peaks, the enhancer-only model showed high correlation of feature importance metrics between the random forest and ridge regression models (**Figure 2E**). Additionally, we trained the enhancer-only model on different BAF^inh^ timepoints and doses and found high agreement among ridge coefficients and MDI scores (**Supplemental Figure 2H, I**). To characterize features associated with enhancers BAF-dependence and -independence, we ranked all features by ridge coefficient and surveyed: (1) the top 15 features with the largest ridge coefficients, corresponding to features that best predict BAF-independence; (2) the bottom 15 features with the most negative ridge coefficients, corresponding to features that best predict BAF-dependence. Strikingly, four of the top fifteen BAF-dependence features corresponded to members of the Activator Protein 1 (AP-1) transcription factor family (**Figure 2F**). Indeed, we found that enhancers bound by AP-1 transcription factors were significantly more BAF-dependent than other enhancers (**Figure 2G**). The role of AP-1 factors at BAF dependent enhancers is supported by previous studies showing that AP-1 factors physically interact with BAF subunits and can influence BAF recruitment and function [28–30]. In addition to AP-1 family members, highly-ranked features associated with BAF-dependence included transcription factors central to B-cell lineage specification, such as IRF4 [31], POU2F2 [32], and SPI1 (PU.1) [33]. We also note the inclusion of RUNX3, which in addition to roles in B-cell biology is known to represent a specific dependency in Epstein Barr Virus-transformed lines such as GM12878 [34–36]. These findings suggest that binding by tissue-specific factors may be a feature of BAF-dependent enhancers. To quantitatively evaluate this, we calculated tau tissue-specificity indexes (denoted τ below) using GTEx RNA-seq data for all TFs included in our model (**Supplemental Table 3**) [37–39]. We found significantly elevated τ, indicating more tissue-restricted expression, in top BAF-dependence features relative to BAF-independence features (**Figure 2F, Supplemental Fig 2J**).

We next asked whether the characteristics of BAF-dependent enhancers we observed in GM12878 were generalizable to other cell types. We applied our analysis approach to two additional cell lines for which we had ATAC-seq data with and without BAF perturbation as well as extensive ChIP-seq datasets to characterize chromatin features genome-wide: HepG2 hepatocellular carcinoma cells (24h BAF^inh^; 720 TFs, 9 histone modifications; ATAC-seq generated in this study, ChIP-seq data from ENCODE; **Supplemental Tables 4-5**), and H1 embryonic stem cells (ESCs; 24h ACBI1 PROTAC treatment; 62 TFs, 27 histone modifications; **Supplemental Table 6**) [40]. In both cell types, AP-1 TFs were included among the top features associated with BAF-dependence (**Supplemental Figure 2K**). Additionally, features predictive of BAF-dependence in HepG2 were enriched for tissue-specific expression (**Supplemental Figure 2L**). To facilitate direct comparison between cell types, we restricted our analysis to chromatin features that were shared between all three cell types (n = 45) and re-trained models for each. Interestingly, in this analysis we identified some broadly expressed TFs that were consistently associated with BAF-dependence: RE1 Silencing TF (REST; τ = 0.36), which suppresses ectopic expression of neuronal gene programs and positions nucleosomes in a BAF-dependent manner [41, 42] and Yin Yang 1 (YY1, τ = 0.20), which functions in enhancer-promoter looping and has been shown to collaborate with BAF to maintain pluripotency circuits in ESCs [43, 44] (**Figure 2H**) [45]. Our results thus highlight two complementary dimensions of BAF function at enhancers, where lineage-specific TFs (e.g. PU.1 in GM12878, HNF4A in HepG2) as well as a select set of more broadly expressed transcriptional regulators (e.g. AP-1, REST, YY1) are predictive of BAF-dependent chromatin accessibility.

### Primed enhancers are distinctly sensitive to BAF inhibition

When examining enhancer features associated with BAF-independence (i.e., retention of chromatin accessibility after BAF^inh^), we were interested to note the presence of H3K27ac and H3K9ac, histone modifications associated with active enhancers (**Figure 2F)**. This suggested to us that BAF might play different roles in maintaining accessibility at enhancers in different activity states. “Active” and “primed” enhancers can be epigenomically defined based on their histone modification states. Active enhancers are dually modified with H3K4me1 and H3K27ac, whereas primed enhancers possess only H3K4me1 (**Figure 3A**) [20].

**Figure 3:**
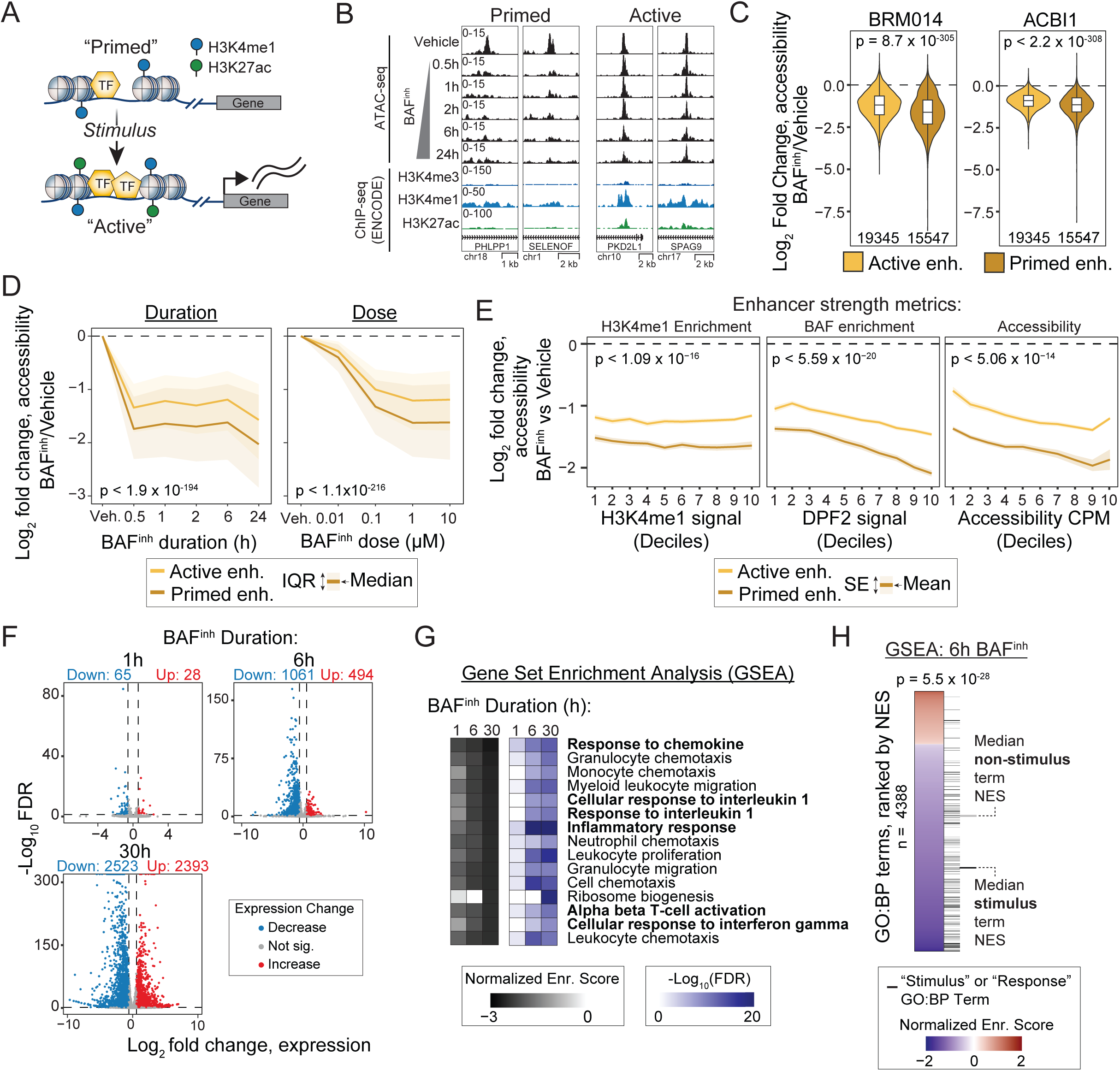
Primed enhancer BAF-dependence and transcriptional changes implicate BAF in stimulus response. (A) Primed enhancers are epigenomically defined by H3K4 monomethylation in the absence of H3K27 acetylation and are activatable upon stimulation. (B) Genome browser examples illustrating more pronounced loss of chromatin accessibility at primed enhancers as compared to active enhancers following BAF^inh^. (C) Primed enhancers show a significantly greater decrease in accessibility compared to active enhancers following BAF perturbation using either BRM014 (6h, 10µM) or ACBI1 (1µM, 24h). P values were calculated using Wilcoxon rank-sum tests; numbers of primed and active enhancers are indicated. (D) Primed enhancers lose accessibility more rapidly (left) and at lower doses (right) than active enhancers. P-values were calculated for each condition using a Wilcoxon rank-sum test. To provide a conservative estimate of significance, the maximum (i.e., least significant) Bonferroni-corrected p-value is reported. (E) Primed enhancers are significantly more BAF-dependent than active enhancers after controlling for three measures of cRE strength: H3K4me1 enrichment (left), enrichment for DPF2, an obligate BAF subunit (middle), and accessibility levels prior to BAF^inh^. Enhancers were binned into deciles for each metric, and P-values were calculated for each decile by a Wilcoxon rank-sum test, and the maximum Bonferroni-corrected p-value is reported. (F) Transcriptional changes following BAF^inh^ (10µM) are biased towards repression at early timepoints (1h, 6h), with more up- and down-regulation at later timepoints. Numbers of genes with significantly increased (FDR < 0.05, log2FC > log2(1.5)) and decreased (FDR < 0.05, log2FC < -log2(1.5)) are indicated. (G) Gene set enrichment analysis (GSEA) reveals consistent downregulation of stimulus-response pathways following BAF inhibition. Heatmaps display normalized enrichment scores (NES; left) and corresponding adjusted p-values (right) for Gene Ontology: Biological Process (GO:BP) terms across all timepoints. The 15 terms shown were selected based on having the most negative mean NES across timepoints. (H) Genes associated with stimulus–response pathways are downregulated following BAF^inh^. GO:BP terms were ranked by Normalized Enrichment Scores (NES) from GSEA (6h BAF^inh^). NES values are displayed as a heatmap, and terms containing “stimulus” or “response” are indicated by black lines to the right of the heatmap. The positions of the median-ranked terms for the “stimulus”/“response” term set and for all other terms are indicated.

To test whether these enhancer classes were differentially BAF-dependent, we first defined GM12878 enhancers based on overlap with H3K4me1 ChIP-seq peaks and subcategorized these into active and primed sets based on H3K27ac peak overlaps (**Supplemental Figure 1J**, **Supplemental Figure 3A**). We found that primed enhancers were broadly BAF-dependent, with 75% of primed enhancers significantly losing accessibility at 30 minutes of BAF inhibition (**Figure 3B, Supplemental Figure 3B**). On average, primed enhancers lost significantly more accessibility than active enhancers following treatment with either BRM014 or the SMARCA2/A4 degrader ACBI1 (**Figure 3C**). Moreover, primed enhancers lost accessibility more rapidly and at lower doses of BAF inhibition than active enhancers (**Figure 3D**). The increased dependence of primed enhancers on BAF to maintain chromatin accessibility is particularly interesting because these enhancers are thought to preconfigure transcriptional responses to stimuli [2, 23, 46–48]. Thus, our finding that primed enhancers are exceptionally BAF-dependent suggests that BAF loss may “decommission” primed enhancers by making them less accessible and thus less able direct stimulus-dependent transcriptional changes.

Before examining stimulus response further, we sought to confirm that the differences in BAF-dependence between primed and active enhancer were truly due to activity status (i.e. primed vs active), rather than other potentially confounding enhancer properties. While primed and active enhancers are distinguished by the presence/absence of H3K27ac, active enhancers also tend to show stronger signal for enhancer-associated features such as H3K4me1 enrichment and chromatin accessibility relative to primed enhancers. Thus, to confirm that the differences in BAF-dependence we observed were driven by activity status rather than by enhancer-associated feature strength, we evaluated the relationship between BAF-dependence and levels of H3K4me1 enrichment and baseline level of accessibility (as measured by ATAC-seq in the absence of BAF^inh^). We also evaluated the relationship between BAF-dependence and binding signal for DPF2, a subunit of BAF complexes. After controlling for these features, we consistently observed greater accessibility loss in primed enhancers versus active (**Figure 3E**). Thus, the differential dependence on BAF between active and primed enhancers is not purely a function of cRE strength – or even BAF binding. We note that prior studies have shown that super enhancers (also known as spread enhancers) are less sensitive to loss of BAF activity than other enhancers [19, 49]. Consistent with this, we found that super-enhancers were even less sensitive to BAF inhibition than typical active enhancers in GM12878 (**Supplemental Figure 3C**).

### Stimulus-associated gene sets are repressed with BAF inhibition

We next asked whether the pronounced changes in enhancer accessibility that we observed in response to BAF^inh^ were accompanied by transcriptional dysregulation. We treated GM12878 cells with BAF^inh^ for 2h, 6h, and 30h and performed RNA sequencing (RNA-seq). Previous studies using BAF subunit (SMARCA4) knockout or BAF inhibition have shown that acute BAF inactivation primarily represses transcription, while longer perturbation timepoints show more balanced up- and downregulated genes due to indirect effects and/or compensatory mechanisms [16–18, 50]. Consistent with these findings, we observed predominantly repressed genes at 1h and 6h, with more upregulated genes emerging by 30h (**Figure 3F; Supplemental Table 7**). To identify pathways affected by BAF^inh^ we performed Gene Set Enrichment Analysis (GSEA). Given the established role of primed enhancers in stimulus responses, we were interested to see multiple response-related Gene Ontology Biological Process (GO:BP) terms among the most strongly downregulated pathways (**Figure 3G**). Notably, the most downregulated pathways also included terms that, while not explicitly annotated as response processes, are plausibly downstream of stimulus signaling, such as chemokine- and migration-related pathways. We expanded on this observation by comparing all GO:BP terms containing “stimulus” or “response” with other terms and found that stimulus/response terms were consistently biased towards downregulation, as reflected by significantly more negative GSEA normalized enrichment scores (**Figure 3H**, **Supplemental Figure 3D**). This indicates that BAF inactivation broadly reduces basal expression of response pathways. We observed a similar result in HepG2 cells treated for 24h with BAF^inh^ (**Supplemental Figure 3E, F; Supplemental Table 8**). Notably, we observed stronger statistical evidence for enrichment at early (1h, 6h) compared to late (30h; 24h HepG2) timepoints, suggesting that downregulation of these pathways is associated with direct effects of BAF^inh^. Together with our finding that most primed enhancers are decommissioned upon BAF inhibition, these results further implicate BAF as a critical regulator of transcriptional stimulus responses. Our data suggests that BAF activity is required to maintain both chromatin and transcriptional programs that define a cell’s competence to respond to stimuli.

### BAF activity is acutely required for transcriptional responses to stimuli

To test whether BAF is required for stimulus response, we first sought to identify stimuli that induce robust transcriptional responses in GM12878. Guided by existing literature, and receptor expression levels in GM12878, we selected six candidate stimuli: dexamethasone (Dex) [51, 52], interferon gamma (IFN-γ) [53], interleukin 2 (IL-2) [54], lipopolysaccharide (LPS) [53], tumor necrosis factor alpha (TNF-*α*) [53], and doxorubicin (Dox) [55]. To determine whether these stimuli induced transcriptional responses in GM12878, we treated cells with each stimulus for 6 hours and performed RNA-seq. We found that IFN-γ and Dex robustly induced expression changes, while the other stimuli had minimal effects (**Figure 4A**). Importantly, the transcriptional programs induced by Dex and IFN-γ are largely orthogonal. Of 500 genes induced by either Dex or IFN-γ, only 10 were significantly induced by both, and the transcription fold changes are not correlated (r = -0.197; **Figure 4B**). We thus selected these two stimuli for further investigation.

**Figure 4:**
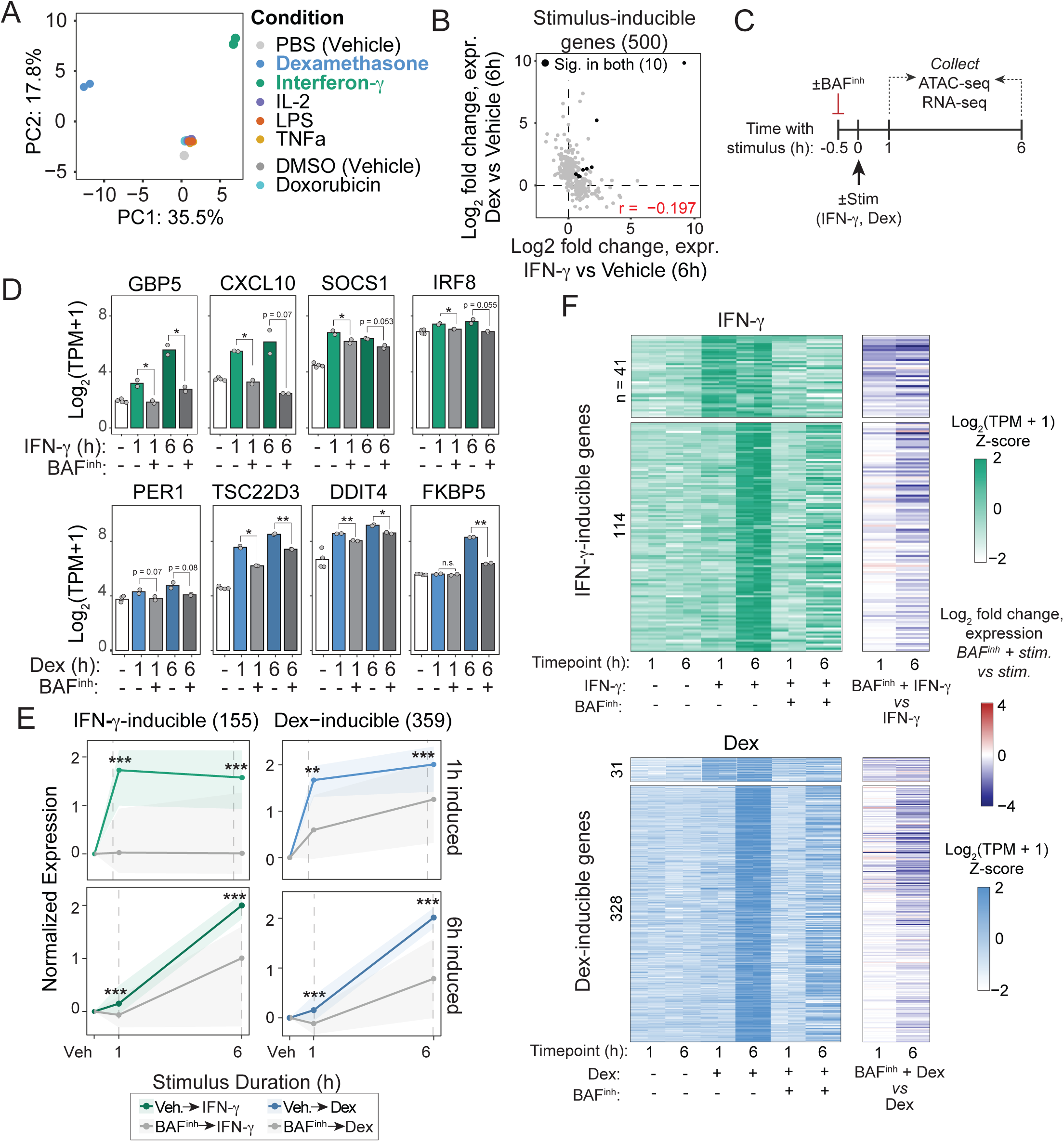
BAF activity is required for full transcriptional responses to interferon gamma (IFN-γ) and dexamethasone (Dex). (A) Global transcriptional responses to candidate stimuli in GM12878, represented by principal component analysis of RNA-seq expression profiles. (B) Transcriptional responses to IFN-γ and Dex are largely distinct. Expression log2FCs for all genes inducible by either stimulus are plotted, and the number of genes significantly induced (FDR < 0.05, log2FC > log2(1.5)) in both conditions is indicated. (C) Experimental approach. Cells were treated with either IFN-γ or Dex, with or without a 30-min pretreatment with BAF^inh^ (10µM). Cells were collected for ATAC-seq and RNA-seq at 1h and 6h post-stimulus. (D) Induction of canonical IFN-γ and Dex targets is suppressed by BAF^inh^ pretreatment. P-values were calculated using a Student’s *t*-test. (E) Both 1h- and 6h-induced genes show attenuated induction with BAF^inh^ pretreatment. Expression values are Z-scored log2(TPM+1), normalized to vehicle control. P-values were calculated using a Wilcoxon rank-sum test. (F) For each stimulus-induced gene, normalized expression levels and log2 fold-change values comparing BAF^inh^ + stimulus versus stimulus alone are shown, illustrating a global reduction in transcriptional induction upon BAF inhibition. *, p < 0.05; **, p < 0.01; ***, p < 0.001, n.s., not significant.

IFN-γ and glucocorticoid signaling drive multi-phasic transcriptional responses, comprising rapid primary induction within minutes to an hour followed by secondary waves of transcription with prolonged stimulation [56, 57]. To capture both early and late aspects of the responses to these stimuli, we profiled transcriptional responses after 1h of treatment in addition to the 6h timepoint previously assessed. As expected, we observed a limited set of transcripts upregulated in each stimulus after 1h (41 and 31 for IFN-γ and Dex, respectively), with broader transcriptional induction at 6h (139 and 355 genes upregulated, respectively) (**Supplemental Figure 4A; Supplemental Tables 9-10**). Dex treatment also resulted in substantial transcriptional repression (15 and 287 genes at 1h and 6h, respectively), consistent with the well-established immunosuppressive effects of glucocorticoids [58, 59]. To validate stimulus-specific transcriptional induction by Dex and IFN-γ, we examined a panel of well-established inducible genes for each treatment. IFN-γ stimulation consistently induced GBP5 [60], CXCL10 [61], SOCS1 [62], and IRF8 [63], whereas Dex treatment upregulated PER1 [64], TSC22D3 [65–67], DDIT4 [65, 66], and FKBP5 [65] (**Supplemental Figure 4B**). Consistent with these gene-level responses, GSEA revealed significant enrichment of “Response to Interferon Gamma” and “Cellular Response to Glucocorticoid Stimulus” biological process terms in IFN-γ– and Dex-treated conditions, respectively (**Supplemental Figure 4C**). For both Dex and IFN-γ, we identified two categories of genes: those that were upregulated at 1h and remained highly expressed at 6h, and those that were induced at 6h only (**Supplemental Figure 4D, E**). Thus, these stimuli provide an ideal pair to probe a generalizable role for BAF in stimulus response (i.e. not specific to a given pathway).

We next tested the requirement for BAF in these transcriptional programs. We pretreated cells with 10µM BAF inhibitor for 30 minutes prior to stimulation with either Dex or IFN-γ for 1 or 6h, then performed RNA-seq and ATAC-seq and (**Figure 4C**). At this time point, chromatin accessibility is already near maximally reduced (**Figure 1B, C**), whereas transcriptional changes remain minimal at the earliest assayed RNA-seq time point (1 hour; **Supplemental Figure 4A**). Thus, this brief pretreatment duration increases the probability that changes in stimulus response after BAF inhibition are the direct result of chromatin accessibility loss, rather than secondary to subsequent transcriptional changes. Remarkably, all genes in our candidate panels for IFN-γ and Dex showed abrogated induction with BAF^inh^ pretreatment (**Figure 4D**). We then analyzed the effects of BAF^inh^ on all IFN-γ or Dex-inducible genes more broadly. For both stimuli, we saw that the 1h-induced and 6h-induced gene sets largely failed to induce (**Figure 4E, F; Supplemental Tables 9-10**). Overall, a majority of inducible genes showed significantly reduced expression with BAF^inh^ pretreatment + stimulus compared to stimulus alone (FDR < 0.05, log2FC < -log2(1.5); 87/155 IFN-γ inducible, 183/359 Dex-inducible). Moreover, most remaining genes showed reduction in expression despite not meeting gene-level significance thresholds (58/68 IFN-γ, 85%; 139/176 Dex, 79%). Thus, our results clearly demonstrate that the transcriptional responses to Dex and IFN-γ, two distinct biological stimuli, depend on continuous BAF activity.

### Stimulus-responsive chromatin relies on BAF activity for accessibility maintenance

We next evaluated the role of BAF in the chromatin response to these stimuli. Treatment with IFN-γ or Dex led to thousands of differentially accessible chromatin loci, with most changes reflecting gained accessibility (**Supplemental Figure 5A, B; Supplemental Tables 11-12**), In total, we identified 2091 and 3828 regions that significantly gained accessibility with IFN-γ treatment and Dex treatment, respectively (FDR < 0.05, log2FC > log2(1.5)). Paralleling what we observed for gene expression, the accessibility gains we observed for the two stimuli showed minimal overlap (**Supplemental Figure 5C**). Only 98 out of 5821 total inducible regions significantly gained accessibility in both conditions, and we saw no evidence of positive correlation between the two conditions. The transcriptional responses to IFN-γ and Dex are known to depend on the TFs STAT1 and NR3C1 (a.k.a. glucocorticoid receptor), respectively. We therefore evaluated enrichment for STAT1 and NR3C1 motifs in both IFN-γ- and Dex-inducible regions and observed stimulus-specific enrichment (IFN-γ: 2.71-fold; Dex: 6.5-fold), supporting the relevance of these loci (**Supplemental Figure 5D**). Given the established role of enhancers in mediating signal-responsive transcription, we asked whether gained-accessibility regions were enriched for enhancer elements. Indeed, we saw strong enrichment of enhancers among induced regions in both conditions (IFN-γ: OR = 6.96, p = 7.37 x 10^-139^; Dex: OR = 6.57, p = 6.33 x 10^-283^, Fisher’s exact test; **Supplemental Figure 5E**), suggesting that enhancers play a major role in driving the transcriptional responses to both stimuli. Despite similarly robust overall accessibility gains, the two stimuli differed markedly in their temporal dynamics. Most IFN-γ-induced regions were specific to a single timepoint, with only 142 loci significantly induced at both 1 h and 6 h (**Supplemental Figure 5F, G**). In contrast, nearly half of Dex-inducible regions (1,691/3,828) were shared across timepoints, and regions significant at only one Dex timepoint typically showed concordant, though sub-significant, gains at the other.

To further assess the involvement of these cREs in stimulus-responsive transcription, we leveraged published GM12878 Promoter Capture Hi-C (PCHi-C) to identify physical contacts between genes and distal cREs [68]. As both primed and active enhancers engage in chromatin looping [69], PCHi-C should capture both active and potentially stimulus-inducible enhancer–gene contacts. We first asked whether stimulus-inducible genes tend to be linked by PCHi-C loops to cREs that gain accessibility upon stimulus. Indeed, we observed significant enrichment for PCHi-C links between inducible genes and inducible cREs for both IFN-γ (OR = 2.58, p = 7.11 x 10^-21^, Fisher’s exact test) and Dex (OR = 2.38, p = 6.40 x 10^-84^). In contrast, this enrichment was largely absent when considering mismatched stimulus pairs (i.e. Dex gene-IFN-γ cRE: OR = 1.19; IFN-γ gene-Dex cRE: OR = 1.21). We next asked, conversely, whether stimulus-inducible cREs tend to be linked to genes that gain expression upon stimulus. We found that genes linked to stimulus-inducible enhancers exhibited stimulus-dependent expression gains, with significant effects extending to interactions of >100Kb (**Supplemental Figure 5H**). Again, significant effects were observed only for matched stimulus conditions. Taken together, these results demonstrate that IFN-γ and Dex induce pronounced, enhancer-driven, and stimulus-specific chromatin and transcriptional responses.

Having characterized stimulus-induced chromatin accessibility changes, we next asked whether these changes depend on BAF chromatin remodeling activity (**Figure 5A).** We found that BAF^inh^ pretreatment abrogated stimulation-associated accessibility gains at both IFN-γ- and Dex-inducible regions (**Figure 5B, C; Supplemental Tables 11-12**). This failure to gain accessibility was observed for both early (1h) and late (6h)-induced regions (**Figure 5D)**. Notably, while some regions retained basal accessibility following BAF^inh^, many loci instead exhibited near-complete loss of accessibility even prior to stimulation (**Figure 5B**). To distinguish this more extreme “decommissioning” phenomenon (i.e. losing accessibility prior to stimulation) from a more subtle failure to induce (i.e. failure to gain further accessibility upon stimulation), we stratified regions based on whether BAF^inh^ alone resulted in significant accessibility loss. Notably, both classes showed impairment of stimulus-induced accessibility gains, indicating that even regions not fully dependent on BAF for basal accessibility nonetheless require BAF activity for proper chromatin remodeling in response to stimulation (**Supplemental Figure 5I**). Our results establish an acute and continuous requirement for BAF activity in chromatin responses to distinct stimuli in GM12878.

**Figure 5:**
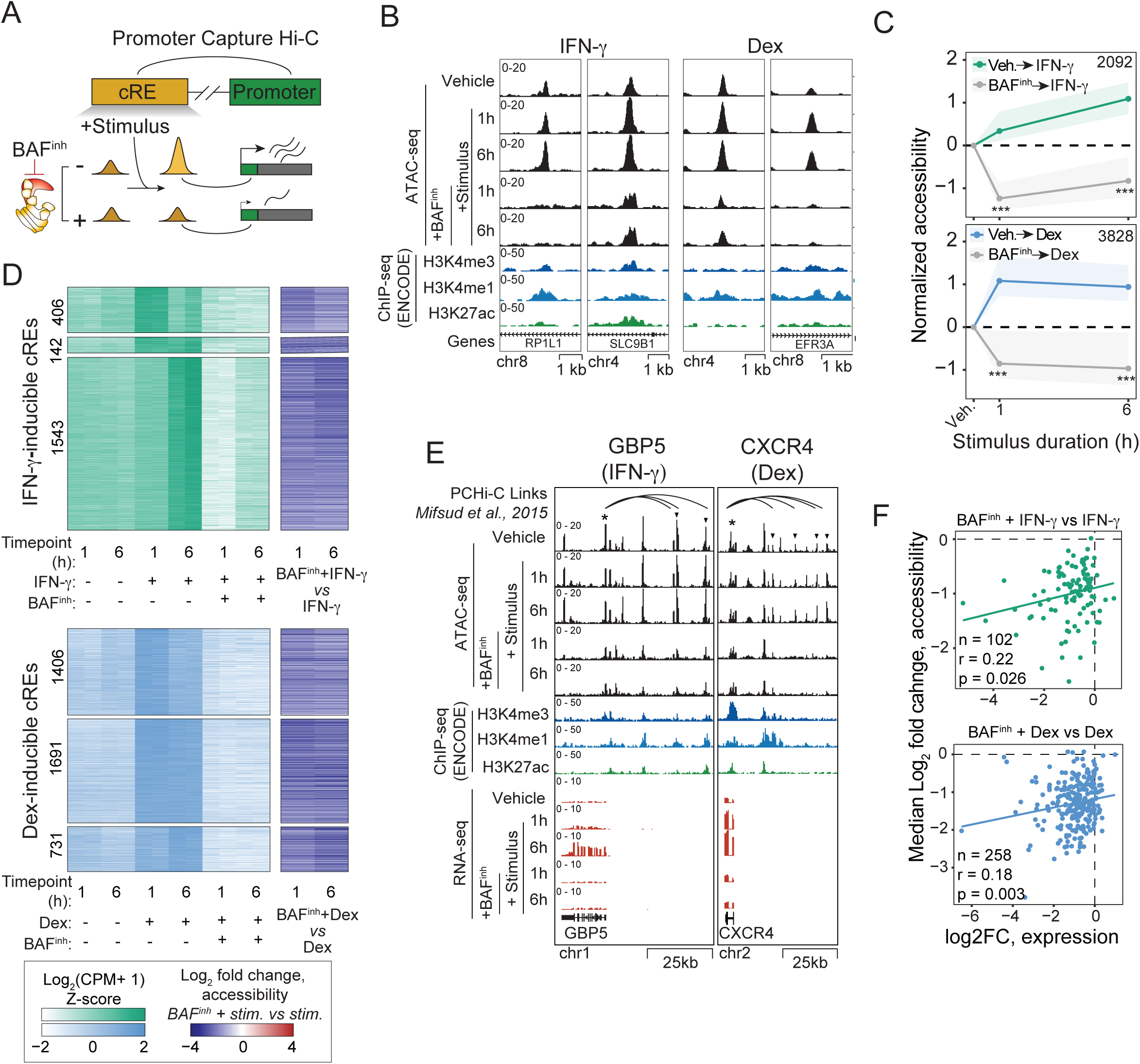
BAF^inh^ restrains stimulus-responsive chromatin accessibility gains and is associated with impaired transcriptional induction at linked genes. (A) Experimental design. GM12878 Promoter Capture Hi-C contacts [68] were used to link IFN-γ and Dex-inducible genes with distal cREs, enabling association of defects in stimulus-responsive chromatin accessibility gains following BAF^inh^ pretreatment to impaired transcriptional upregulation. (B) Genome browser examples of enhancers that gain accessibility with IFN-γ (left) or Dex (right) treatment but fail to be induced with BAF^inh^ pretreatment. (C) IFN-γ- and Dex-induced accessibility gains fail to occur with BAF^inh^ pretreatment. Accessibility is shown as Z-scored Log2(CPM+1), normalized to vehicle. Shaded regions represent the interquartile range. P-values were calculated using Wilcoxon rank-sum tests (***, p < 0.001). (D) For each IFN-γ (top) or Dex (bottom) cRE, normalized accessibility levels and log2 fold-change values comparing BAF^inh^ + stimulus versus stimulus alone are shown. cREs are grouped based on significant induction at 1 h only (top), both time points (middle), or 6 h only (bottom); numbers of elements in each category are indicated. (E) Genome browser examples of stimulus-induced, BAF-dependent genes with promoter capture Hi-C (PCHi-C)–linked cREs indicated by arcs. Asterisks denote promoters and arrowheads indicate BAF-dependent linked cREs. (F) Reductions in stimulus-dependent accessibility gains are associated with reduced transcriptional induction at physically linked genes.

We next revisited the PCHi-C gene-cRE interactions to ask whether BAF-dependent stimulus-responsive cREs were linked to genes that failed to induce upon stimulation (**Figure 5A**). We first observed that canonical response genes for IFN-γ (e.g, GBP5) and Dex (e.g., CXCR4), which showed defective induction with BAF^inh^ pretreatment, made multiple PCHi-C contacts with BAF-dependent cREs (**Figure 5E**). We next expanded our analysis to include all inducible genes for which promoter capture Hi-C (PCHi-C) contacts were available (IFN-γ: 102/155; dexamethasone: 258/359). Within these sets, we focused on inducible genes whose transcriptional induction was significantly reduced by BAF inhibition (59 IFN-γ–responsive genes; 143 Dex-responsive genes). Notably, all such genes (59/59 IFN-γ; 143/143 Dex) were linked by PCHi-C to at least one candidate regulatory element (cRE) that exhibited reduced accessibility following BAF^inh^ pretreatment. Moreover, many of these genes were linked to at least one cRE that normally gained accessibility upon stimulation but failed to do so when pretreated with BAF^inh^ (29/59 IFN-γ; 108/143 Dex). As genes are often linked with multiple cREs, we aggregated the effect of BAF^inh^ on cRE accessibility on a per-gene basis by taking the median change in accessibility across all linked elements. For both stimuli, we found that reduced transcriptional induction was significantly although modestly correlated with reduced chromatin accessibility gains (**Figure 5F**). In summary, we find that BAF-dependent, stimulus-responsive enhancers are linked to BAF-dependent, stimulus-responsive genes, revealing dual roles for BAF in maintaining enhancer accessibility and enabling its dynamic modulation upon stimulation.

## DISCUSSION

In this study we systematically evaluated the role of BAF in shaping the accessibility and stimulus-responsiveness of regulatory chromatin. In GM12878 cells, we uncovered a distinct reliance of primed enhancers on BAF activity and extended this observation to show a broad requirement for BAF at stimulus-responsive chromatin. Disruption of BAF activity suppresses both transcriptional and chromatin responses to two distinct stimuli, interferon gamma and dexamethasone, highlighting the critical role of this complex in signal response. Our results support a model where maintenance of stimulus-responsive enhancer accessibility by BAF licenses subsequent enhancer activation and transcriptional induction (**Figure 6A**). Thus, inhibition of BAF compromises stimulus responses by preventing enhancer activation (**Figure 6B**). Given that heterozygous putative loss-of-function BAF subunit mutations are recurrently observed in disease [7, 70], it is intriguing to speculate that impaired stimulus responsiveness may contribute to aberrant phenotypes by altering cell differentiation, cellular function, or both.

**Figure 6:**
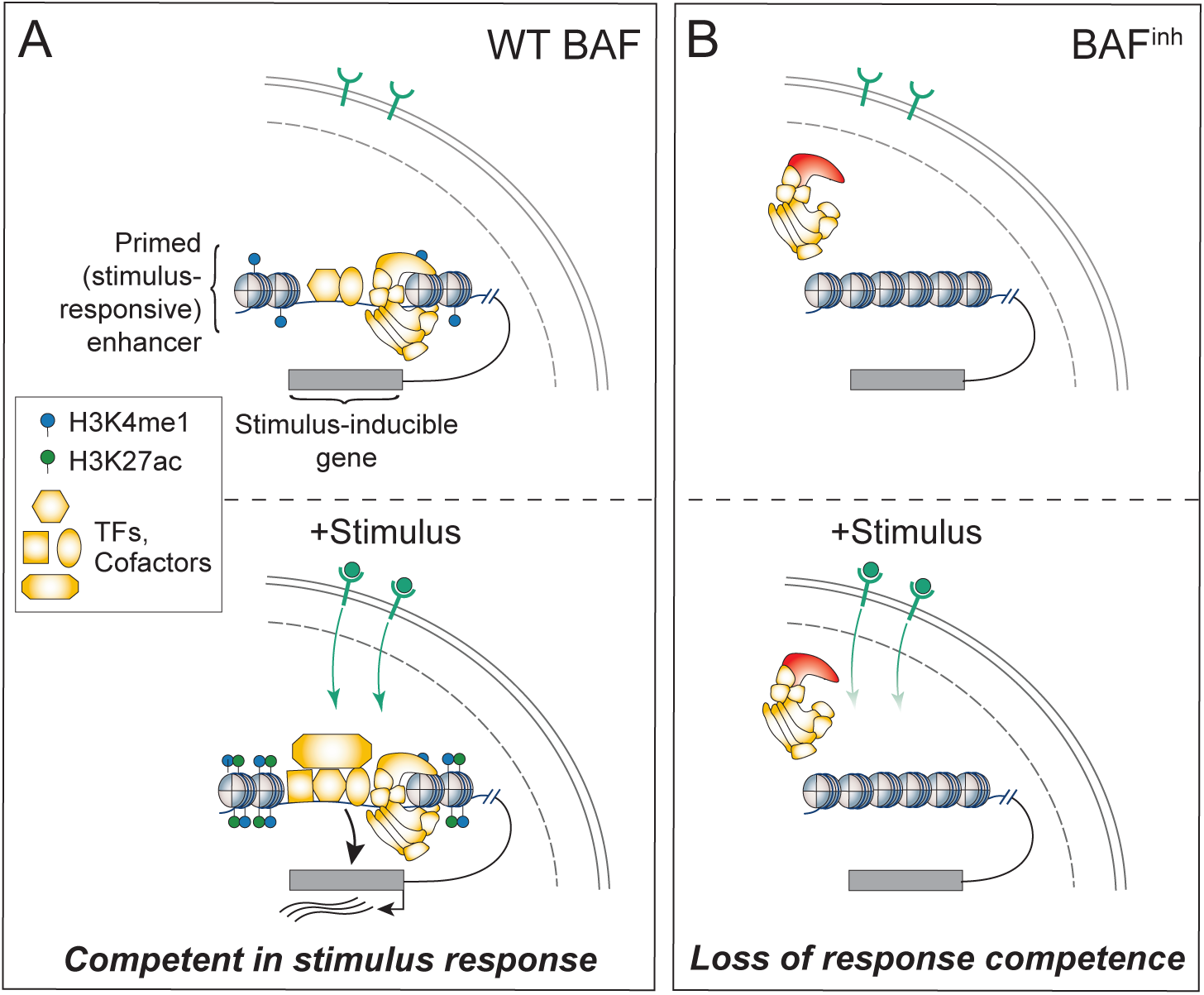
Model for BAF regulation of stimulus-responsive transcription through maintenance of primed enhancer accessibility. (A) BAF complexes continuously maintain chromatin accessibility at primed enhancers. Upon stimulation, signal-dependent TF activity promotes enhancer activation and transcriptional induction of target genes (bottom). (B) Reduction of BAF activity leads to widespread loss of accessibility at primed enhancers (top), impairing enhancer activation and resulting in defective transcriptional responses to stimuli (bottom).

Genetic or pharmacological perturbations of BAF activity have shown that classes of cREs vary in their BAF-dependence, with enhancers exhibiting greater dependence and promoters and CTCF sites remaining mostly unaffected [15, 16, 18, 19, 49]. However, the basis for within-class heterogeneity -particularly within enhancers – has remained poorly understood. We addressed this gap by developing a machine learning framework to model enhancer-specific accessibility changes and predict BAF dependence. Using this approach, we identified features including AP-1 TFs and lineage-determining factors that predict BAF-dependence. Unlike prior approaches that modeled accessibility changes across all regulatory elements [71] or used deep learning to identify predictive DNA sequence features [19], our strategy leverages ChIP-seq features to identify individual transcription factor family members associated with enhancer-specific BAF dependence. By identifying the molecular features that confer sensitivity to BAF inhibition at enhancers, our work provides a foundation for understanding how BAF loss of function affects the regulatory landscape in disease. Importantly, the machine learning framework we developed has the potential for broad applicability (beyond BAF), as it can be applied to identify features predictive of accessibility changes following any perturbation.

Our finding that AP-1 TF binding predicts BAF-dependence, and that AP-1-bound enhancers are distinctly BAF-dependent, expands on a growing body of evidence linking BAF with this TF family. AP-1 TFs physically interact with BAF complex subunits and have been implicated in the recruitment of this complex [28, 30]. Across diverse cell contexts, AP-1 TF binding and motif enrichment is enriched at regulatory elements that lose accessibility or activity (as measured by H3K27ac) following BAF perturbation [49, 72–74]. Together, our results and prior studies, along with the ubiquitous expression of AP-1 TFs, point towards a general role of AP-1 in BAF recruitment and/or function. Interestingly, AP-1 TFs are involved in a variety of stimulus responses and have known roles in enhancer activation in response to mitogenic stimuli, cytokines, and glucocorticoids [30, 51, 75]. In these contexts, disruption of AP-1 binding results in attenuation of stimulus-dependent chromatin accessibility changes. This complements our observations that both epigenomically-defined primed enhancers (H3K27ac-H3K4me1+) and enhancers linked to stimulus-responsive genes are distinctly sensitive to BAF inhibition. Together, this raises the interesting possibility that BAF and AP-1 cooperate to maintain stimulus-responsive chromatin. Notably, however, AP-1 TFs are not thought to play a direct role in the IFN-γ response [76]. Thus, our observation that BAF inhibition strongly suppresses the response to this molecule indicates that BAF’s contribution to stimulus-dependent chromatin modulation extends beyond AP-1 regulatory contexts.

The present study also builds on a substantial body of prior work linking BAF to a variety of stimulus responses. The adrenal carcinoma cell line SW-13, which lacks SMARCA4 expression, exhibits defective transcriptional responses to diverse stimuli, including type I and II interferons as well as steroid hormone signaling [77–83]. Additionally, depletion of SMARCA2/A4 or inhibition of BAF ATPase activity has been shown to impair responses to neuronal depolarization, glucocorticoids, hypoxia, Notch signaling, and the bacterial endotoxin lipopolysaccharide/lipid A [84–90]. Most recently, studies of lipid A and Notch signaling have indicated that BAF activity is *acutely* required for stimulus-induced transcription [87, 88]. Within this framework, our study makes two major advances. First, by acutely inhibiting BAF activity and jointly profiling chromatin accessibility and transcription, we provide a more temporally resolved understanding of BAF function in IFN-γ and dexamethasone responses and define the extent to which these responses depend on continuous BAF activity. Second, by establishing IFN-γ and dexamethasone as additional stimulus contexts in which ongoing BAF activity is required to enable chromatin remodeling and transcriptional induction, our findings strengthen the case for a broad role for BAF complexes in stimulus-responsive gene regulation.

Although our use of orthogonal stimuli and the extensive prior literature on BAF in stimulus response suggest a broad role for BAF in regulating inducible chromatin, the full generalizability of our findings remains unclear. In particular, the extent to which BAF-dependence of primed and stimulus-responsive enhancers extends to other signaling pathways, cell types, or developmental contexts remains to be determined. Notably, while stimulus-inducible chromatin accessibility was overwhelmingly BAF dependent, some stimulus-inducible genes are able to increase transcription at least partially following BAF inhibition. Understanding the compensatory mechanisms that enable BAF-independent transcriptional induction may provide insight into the regulatory logic of stimulus-responsive gene expression.

## CONCLUSIONS

Collectively, our work highlights the role of BAF complexes as central regulators of stimulus-responsive chromatin accessibility and transcription. In addition to identifying a continuous requirement for BAF activity in maintaining primed and stimulus-inducible enhancers, this study establishes experimental and computational frameworks for investigating the effects of perturbations on chromatin remodeling in other contexts. Future studies can build on our findings to define the functional interplay between BAF and stimulus-associated transcription factors, and to assess the broader contribution of BAF in shaping cellular responsiveness to signaling cues in healthy as well as disease states.

## METHODS

### Cell Culture

GM12878 cells were cultured in suspension using RPMI-1640 medium (Thermo Scientific, A1049101) supplemented with 2 mM L-glutamine (Thermo Scientific, 25030081), 15% fetal bovine serum (FBS; Fisher Scientific, 10-082-147), and 100U/mL penicillin/streptomycin (Fisher Scientific, 15-140-122). HepG2 cells were cultured in Eagle’s Minimal Essential Medium (ATCC, 30-2003) supplemented with 10% FBS and 100U/mL penicillin/streptomycin. HepG2 cells were passaged using Trypsin-EDTA (Fisher Scientific, 25-200-072). All cells were maintained at 37 °C in a humidified incubator with 5% CO₂. Viability was measured using Trypan Blue staining (Millipore Sigma, T8154).

### BAF^inh^ and Stimulus Treatment

BRM014 (MedChemExpress, HY-119374) was prepared as a 10mM stock in dimethyl sulfoxide (DMSO) and used at a final concentration of 10µM unless otherwise indicated. Vehicle controls were treated with an equivalent concentration of DMSO. ACBI1 (opnMe, ACBI1) was similarly prepared as a 10mM stock in DMSO and used at a final concentration of 1µM unless otherwise indicated.

For the initial transcriptional stimulus panel in Figure 4A, concentrations were determined based on literature review. Briefly, cells were treated with lipopolysaccharide (Sigma L9641, 10ng/mL)[53, 91], tumor necrosis factor alpha (Genscript Z01001, 25ng/mL) [92, 93], interferon gamma (Genscript, Z02986, 100ng/mL) [53], interleukin 2 (Sigma, 11011456001, 50U/mL) [54], dexamethasone (MedChemExpress, HY-14648, 500nM) [51, 52], or doxorubicin (Selleck Chem, E2516, 500nM) [55, 94] for 6h. For BAF^inh^ pretreatment, 10µM BRM014 was added to cells 30min prior to the addition of either 100ng/mL IFN-γ or 500nM Dex for 1h or 6h.

### Western Blotting

GM12878 cells were treated for 24 h with ACBI1 (0.1 or 1 µM) or BRM014 (1 or 10 µM). Vehicle (DMSO)–treated and untreated controls were included. Cells were pelleted by centrifugation at 500 x g for 5 min at 4 °C, washed once with ice-cold 1x PBS, and resuspended in ice-cold RIPA buffer (50 mM Tris-HCl pH 7.4, 150 mM NaCl, 0.5% w/v sodium deoxycholate, 0.1% sodium dodecyl sulfate, 1% v/v IGEPAL CA-630) supplemented with 1x cOmplete EDTA-free Protease Inhibitor Cocktail (Millipore Sigma, 11873580001). Lysates were incubated for 30 min at 4 °C and clarified by centrifugation at 14,000 rpm for 10 min at 4 °C.

For immunoblotting, 15 µL of each protein extract was resolved by SDS–PAGE on acrylamide gels run at 160 V and transferred to nitrocellulose membranes using the Trans-Blot Turbo system (1.3 A, 10 min). Protein transfer was confirmed by Ponceau S staining. Membranes were blocked for 30 min at room temperature in 5% non-fat dry milk in 1× PBS containing 0.1% Tween-20 and probed for 2 h at room temperature with anti-SMARCA2 (1:1000; Cell Signaling Technology, 11966), anti-SMARCA4 (1:1000; Cell Signaling Technology, 49360), or anti-GAPDH (1:5000; Cell Signaling Technology, 2118) antibodies. After washing, membranes were incubated for 1h at room temperature with anti-rabbit IgG–HRP secondary antibody (1:2000; Cell Signaling Technology, 7074). Signal was developed using Clarity ECL Substrate (Bio-Rad, 1705060) and imaged on a Bio-Rad ChemiDoc system.

### ATAC-seq

ATAC-seq was performed as previously described [95], with some modifications as described below. All experiments were performed with a minimum of two biological replicates per condition. Cells were harvested by centrifugation at 500 x g for 5 min, resuspended in cold base media (GM12878: RPMI-1640; HepG2: EMEM) supplemented with 10% DMSO, and immediately flash-frozen in liquid nitrogen to minimize recovery of chromatin accessibility. Samples were stored at -80°C until use.

Frozen samples were rapidly thawed in a 37°C water bath and cells were pelleted by centrifugation. After washing with cold 1x PBS, cells were resuspended in cold nuclear permeabilization buffer (1x PBS, 5% w/v bovine serum albumin, 0.2% w/v IGEPAL CA-630, 1mM dithiothreitol, 1x cOmplete EDTA-free Protease Inhibitor Cocktail) and nuclei were extracted by incubation with rocking for 5 minutes at 4°C. Nuclei were pelleted and resuspended in tagmentation buffer (33mM Tris-acetate pH 7.8, 66mM potassium acetate, 11mM magnesium acetate, 16% v/v dimethylformamide) and 50,000 nuclei were tagmented for 1h at 37°C with Tn5 (Illumina, 20034197). Tagmented DNA was purified using the MinElute PCR Purification Kit (Qiagen, 28004) and amplified by 7-8 cycles of PCR using NEBNext High-Fidelity Q5 polymerase (New England Biolabs, M0541). Amplified libraries were size-selected using SPRIselect beads (Beckman Coulter, B23318), quantified using a Qubit fluorometer (Thermo Fisher Scientific) with dsDNA High Sensitivity Assay reagents (Thermo Fisher Scientific, Q32854), and library size distributions was evaluated using High Sensitivity D1000 ScreenTapes on an Agilent 4200 TapeStation system. Libraries were sequenced (150bp paired-end) on an Illumina NovaSeq platform.

### RNA-seq

All experiments were performed with a minimum of two biological replicates per condition. Cells were harvested by centrifugation at 500 x g for 5 min, and after removing supernatant cell pellets were flash frozen in liquid nitrogen and stored at -80°C until use. Cells were disrupted using QIAshredder columns (Qiagen, 79656) and total RNA was prepared using RNeasy Mini kits (Qiagen, 74104) as per manufacturer protocol. RNA quality was assessed using High Sensitivity RNA ScreenTapes (Agilent, 5067-5580) run on an Agilent 4200 TapeStation system. All samples had RINe values greater than 9.0. PolyA-enriched, stranded RNA-seq libraries were generated from 500ng total RNA for each sample using the Illumina Stranded mRNA prep kit (Illumina, 20040534) following manufacturer protocol. Library quality control and sequencing was performed as described for ATAC-seq libraries.

### ATAC-seq Data Analysis

#### Data processing

Quality control on raw FASTQs was performed using FastQC [96] and FastQ Screen [97]. Data were processed using the Encyclopedia Of DNA Elements (ENCODE) ATAC-seq Pipeline, (v2.2.0, https://github.com/ENCODE-DCC/atac-seq-pipeline) [98], executed using Singularity containers. Briefly, adapter trimming was performed using cutadapt [99] with parameters -e 0.1 -m 5 and reads were aligned to the hg38 reference genome using Bowtie2 [100]. Reads were filtered using SAMtools [101], PCR duplicates were flagged and removed using Picard [102], and peaks and signal tracks were generated using MACS2 [103]. For each condition, replicated peak sets were generated using the intersectBed utility from the bedtools suite [104] by retaining peaks present both in the pooled peak set and in all individual replicates. A consensus GM12878 peak set was generated by concatenating and merging replicated DMSO and BRM014 (t = 6h; n = 3 replicates per condition) peaks. Read count matrices were produced from filtered BAM files using the bedtools multiCov tool [104]. Consensus peaks were filtered to exclude low-count peaks (DMSO mean counts < 10), yielding a final set of 111,721 peaks. Applying the same approach to HepG2 (t = 24h, n = 2 vehicle and n = 2 BAF^inh^) and H1 (t = 24h, n = 3 vehicle and n = 3 ACBI1) [40] ATAC-seq datasets yielded 148,873 and 177,081 peaks, respectively.

#### Differential accessibility analysis

The global loss in accessibility resulting from BAF^inh^ precludes use of standard normalization strategies when performing differential accessibility analysis using published tools. We therefore adopted a normalization approach based on prior work [105]. Transcriptomic data were used to identify genes with invariant expression between DMSO and BAF^inh^ conditions (FDR > 0.2 and < 20% change in expression; n = 6,263 genes) and library size factors were calculated for each sample as the sum of ATAC-seq read counts in the promoters of these genes. This strategy assumes that promoter accessibility at invariantly expressed genes reflects sequencing depth rather than biological signal. Differentially accessible regions were identified using the quasi-likelihood framework implemented in edgeR (*glmQLFit* and *glmQLFTest*) [106]. Differentially accessible regions were defined as those with FDR < 0.05 and an absolute log2FC > 1 unless otherwise stated.

#### cRE Annotation

Peaks were assigned cRE type annotations using three different approaches. First, we used a published 15-state ChromHMM model (GEO Accession: GSM936082; [25]). Chromatin state annotations were lifted over from hg19 to hg38, and overlaps between chromatin states and ATAC-seq peaks were identified using the GenomicRanges Bioconductor package [107]. For peaks overlapping multiple ChromHMM states, the state with the largest genomic overlap was assigned. An analogous approach was used to assign annotations from the ENCODE candidate cis-regulatory element (cCRE) catalog [26] to our ATAC-seq peaks. Finally, we used a combination of transcription start site (TSS) coordinates and ENCODE ChIP-seq data to classify ATAC-seq peaks, summarized in **Supplemental Figures 1J and 3A**. GENCODE v47 comprehensive gene annotations were used to define 1 kb windows centered on annotated TSSs. ATAC-seq peaks overlapping these windows were classified as TSS-proximal, and all remaining peaks were classified as TSS-distal. TSS-proximal peaks overlapping H3K4me3 ChIP-seq peaks (ENCODE accession: ENCFF636FWF) were annotated as **promoters**. TSS-distal ATAC-seq peaks overlapping H3K4me1 ChIP-seq peaks (ENCFF453PEP) were further classified as **active enhancers** if they also overlapped H3K27ac (ENCFF361XMX), **poised enhancers** if they overlapped H3K27me3 (ENCFF035PQG), or **primed enhancers** if they lacked overlap with either H3K27 modification. TSS-distal peaks lacking H3K4me1 but overlapping CTCF ChIP-seq peaks (ENCFF827JRI) were annotated as **CTCF-bound**. Super-enhancers were defined using two GM12878 annotations: SEdb v3.0 [108] and dbSUPER [109], comprising 153 and 257 regions, respectively. The union of these two annotation sets was used as our super-enhancer reference. Active enhancers residing within poised enhancers were identified using the Bioconductor GenomicRanges package [107].

#### Transcription Factor Motif Enrichment

Motif position weight matrices for NR3C1 and STAT1 were extracted from the JASPAR2022 database using the TFBSTools Bioconductor package, restricting to human motifs (species ID 9606). Transcription factor motif occurrences within ATAC-seq peaks were identified using the *matchMotifs* function from the Bioconductor motifmatchr package [110]. Motif enrichment was assessed using Fisher’s exact test by comparing the frequency of motif-containing peaks among ATAC-seq peaks that significantly gained accessibility upon stimulation to the background set of all ATAC-seq peaks.

#### Promoter Capture Hi-C

Coordinates and metadata for significant (as defined in [68]) promoter-cRE links from promoter capture Hi-C experiments in GM12878 cells were downloaded from the EMBL-EBI ArrayExpress repository (Accession: E-MTAB-2323). Genomic coordinates were converted to GRanges objects and lifted over from hg19 to hg38 using the *liftOver* function from the rtracklayer package. Genomic distances between promoter and target fragments were calculated using the *distance* function from the GenomicRanges package. ATAC-seq peaks overlapping prey (non-promoter) fragments were identified using the *findOverlaps* function from GenomicRanges.

#### Data visualization

Genomic heatmaps were generated with the *computeMatrix* and *plotHeatmap* utilities from the deepTools suite [111] using fold change over control bigWig files as inputs. bigWig tracks representing BAF^inh^ vs Vehicle log2 fold change in accessibility were generated using the deepTools *bigwigCompare* tool. We used the *bigWigAverageOverBed* command line tool [112] to quantify H3K4me1 and DPF2 signal within ATAC-seq peaks. Peaks and signal tracks were visualized using the UCSC Genome Browser [112]. All other data visualization was performed in R using ggplot2 [113], with heatmaps produced using pheatmap [114].

### RNA-seq Data Analysis

#### Data processing

FastQ quality control was performed as described for ATAC-seq data. Data were processed using the ENCODE RNA-seq pipeline (v1.2.4, https://github.com/ENCODE-DCC/rna-seq-pipeline) [98], executed using Singularity containers. Reads were mapped to the hg38 reference genome using STAR [115] and gene-level expression quantifications were produced using RSEM [116].

#### Differential expression analysis

RSEM outputs were imported to R using tximport [117] and filtered to exclude genes with a maximum TPM of ≤ 1 or a length of 0. Principal component analysis (PCA) was performed on variance-stabilized gene expression values generated with DESeq2 [118] using the prcomp function in R. Differential expression analysis was performed using DESeq2 [118]. We defined differential expression as FDR < 0.05 and | log2FC| ≥ log2(1.5).

#### Gene set enrichment analysis

Gene set enrichment analysis was performed using the fgsea Bioconductor package [119], with genes ranked by log2 fold change from differential expression analysis. The background gene set consisted of all genes expressed (after filtering, as described above) in the relevant cell line (GM12878 or HepG2). Gene sets containing fewer than 10 genes were excluded from analysis.

##### Data visualization

RNA-seq signal tracks were visualized using the UCSC Genome Browser and are publicly viewable (see above); all other visualization was done in R using ggplot2 and pheatmap.

### Random forest and ridge regression modelling

Python code for all Machine-learning (ML) methods outlined in this section is available in the following GitHub repository (https://github.com/aodgulka/accessibility-prediction-from-chipseq/). The code is available in two forms – a series of executable python scripts for end-to-end reproduction and a Jupyter Notebook which outlines step-by-step processes. The former was used for all figures in this study. ML training performance evaluation and feature importance analyses were implemented using the scikit-learn package in Python [120].

For all ML methods, we first generated feature matrices in which each row is an accessible chromatin peak (from consensus, filtered ATAC-seq peaks for each cell type, described above), and each column is ChIP-seq target (histone modification or TF). ChIP-seq datasets were accessed programmatically via the ENCODE API. In cases where individual TFs or histone modifications had multiple experiments available in the ENCODE database, we selected the experiment with the highest Fraction of Reads in Peaks (FriP) score for our analyses. FRiP is often used as a quality metric in ChIP-seq and ATAC-seq experiments [121, 122]. Peak calls (hg38) were downloaded from ENCODE as narrowPeak files, using replicated peak sets for histone ChIP–seq experiments and IDR-thresholded replicated peak sets for TF ChIP–seq experiments. In total, we extracted 166 datasets (155 TF, 11 histone) in GM12878, (720 TF, 9 histone) in HepG2, and 89 (62 TF, 27 histone) in H1. The ENCODE accessions and corresponding FRiP scores for datasets used in this study are included in **Supplemental Table 2**. We used the bedtools intersect tool to generate a binary feature matrix, where 1 indicates overlap between ATAC-seq and ChIP-seq peaks and 0 corresponds to no overlap. This feature matrix was used as the input for our ML approaches.

We trained a Random Forest (RF) Classifier model to identify BAF-dependent (FDR < .05, log2FC < -1) and BAF-independent regions. Prior to model training, the datasets were class balanced (via undersampling) and randomly partitioned into training, validation, and testing sets according to a 70:15:15 ratio. The hyperparameters of the RF were then tuned via a grid search on the validation set, exploring configurations including the number of decision trees (*n_estimators* ∈ *{50, 100, 200, 300}*), maximum tree depth (*max_depth*∈ *{None, 10, 20}*), and minimum samples required per leaf node (*min_samples_leaf* ∈ *{1,2}*). The final hyperparameters of the RF were selected based on the model with the highest mean accuracy. After hyperparameter tuning, we used two metrics, accuracy and Receiver Operating Characteristic (ROC), to evaluate the performance of the trained RF classifier on the test set – the sample of the data which was not seen during training of the model. These metrics are defined as follows:

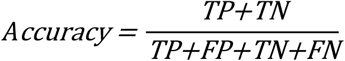

The ROC curve is generated by plotting the True Positive Rate (TPR) against the False Positive Rate (FPR) at different classification thresholds:

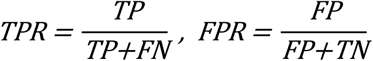

To assess random forest feature importance, we used Mean Decrease in Impurity (MDI) scores as well as permutation-based importance. MDI importance values, which quantify the total reduction in Gini impurity attributable to splits on each feature averaged across all trees, were extracted directly from the fitted model using scikit-learn’s *feature_importances_* attribute. Permutation feature importance was computed on a held-out test set and was calculated via scikit-learn’s *permutation_importance* method with each feature being permuted 10 times.

As a complementary approach, we used ridge-regularized linear regression to predict log2 accessibility fold changes and interpreted the magnitude of fitted coefficients as feature importance and their sign as directionality (a negative coefficient corresponds to BAF-dependence, while a positive coefficient corresponds to BAF-independence). Ridge regression applies an L2-penalty to an ordinary least squared regression model to mitigate multicollinearity in high-dimensional datasets [123]. We chose this over L_1_-penalized, Lasso Regularized Regression, as it enables retention of all features (since coefficients are not pushed to zero) and relative ranking of correlated predictors. In highly correlated datasets with a much larger number of samples compared to the number of features, ridge regression dominates in performance compared to lasso regression [124]. Features were standardized using scikit-learn’s StandardScaler during preprocessing, and the default sk-learn regularization strength (alpha = 1.0) was used during model fitting [120]. To ensure a robust assessment of feature importance, we fitted the ridge regression model 10 times on randomly sampled 70% subsets of the feature matrix (i.e., used bootstrapping) and averaged the coefficients across iterations:

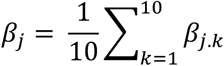

Where *β_j.k_* is the regression coefficient of feature *j* at bootstrapping iteration *k* and *β*_j_ is the final reported ridge regression coefficient for feature *j*.

### Tau index calculation

Tau indexes (τ) are a well-established metric for quantifying tissue-specificity, where scores near 0 and 1 indicate ubiquitous and tissue-restricted expression, respectively [37, 38]. To calculate τ, we utilized expression data from 68 adult tissues available from the Genotype-Tissue Expression (GTEx) portal, v10 release [39]. We generated a gene by tissue expression matrix using median TPM by tissue. After pseudocount adjustment and log2 transformation, τ was calculated for each gene as follows [37]:

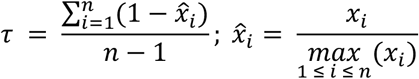

Where *n* is the number of tissues and *x*_*i*_ is the expression of the gene in tissue *i*. We defined tissue-restricted genes as those with τ ≥ 0.8, an established threshold for tissue-specific expression [38]. τ scores for all genes are included in **Supplemental Table 3**.

## DECLARATIONS

### Availability of Data and Materials

Code used to generate the analyses and figures in this study is available at GitHub: https://github.com/GorkinLab/paper-gulka-et-al-baf-inh. ATAC-seq and RNA-seq data generated in this study have been deposited in the Gene Expression Omnibus (GEO) under accessions <u>GSE318082</u> and <u>GSE318083,</u> respectively. H1 ATAC-seq data was downloaded from GEO, Accession: <u>GSE235534</u>. Promoter capture Hi-C data was downloaded from the EMBL-EBI ArrayExpress repository, Accession: <u>E-MTAB-2323</u>.

### Author contributions

The study was conceptualized by AODG and DUG. Experiments were performed by AODG, KAK, and ZZ. Machine learning approach was developed by KAK, AODG, and DUG. Data analysis was performed by AODG and KAK. The manuscript was written by AODG with contributions from all authors.

## Acknowledgements

This study was supported in part by the Emory Integrated Genomics Core (EIGC; RRID:SCR_023529), which is subsidized by the Emory University School of Medicine and is one of the Emory Integrated Core Facilities. Additional support was provided by the Georgia Clinical & Translational Science Alliance of the National Institutes of Health under Award Number UL1TR002378. The content is solely the responsibility of the authors and does not necessarily reflect the official views of the National Institutes of Health.

## SUPPLEMENTAL MATERIALS

**Supplemental Table 1:** Consensus ATAC-seq peaks in GM12878, annotated by cRE type and effect of BAF^inh^.

**Supplemental Table 2:** ENCODE ChIP-seq accessions and FRiP scores for ChIP-seq datasets used as inputs for machine learning in GM12878

**Supplemental Table 3:** Tau indexes calculated from GTEx adult tissue gene expression datasets.

**Supplemental Table 4:** Consensus ATAC-seq peaks in HepG2, annotated by cRE type and effect of BAF^inh^.

**Supplemental Tables 5-6:** ENCODE ChIP-seq accessions and FRiP scores for ChIP-seq datasets used as inputs for machine learning in HepG2 and H1

**Supplemental Tables 7-8:** Effects of BAF^inh^ on gene expression in GM12878 (1h, 6h, 30h) and HepG2 (24h).

**Supplemental Tables 9-10:** Lists of IFN-γ- and Dex-inducible genes, and effects of BAF^inh^ pretreatment on transcriptional induction.

**Supplemental Tables 11-12:** Lists of regions gaining accessibility with IFN-γ and Dex, and effects of BAF^inh^ pretreatment on accessibility increase.

## SUPPLEMENTAL FIGURE LEGENDS

**Supplemental Figure 1.**
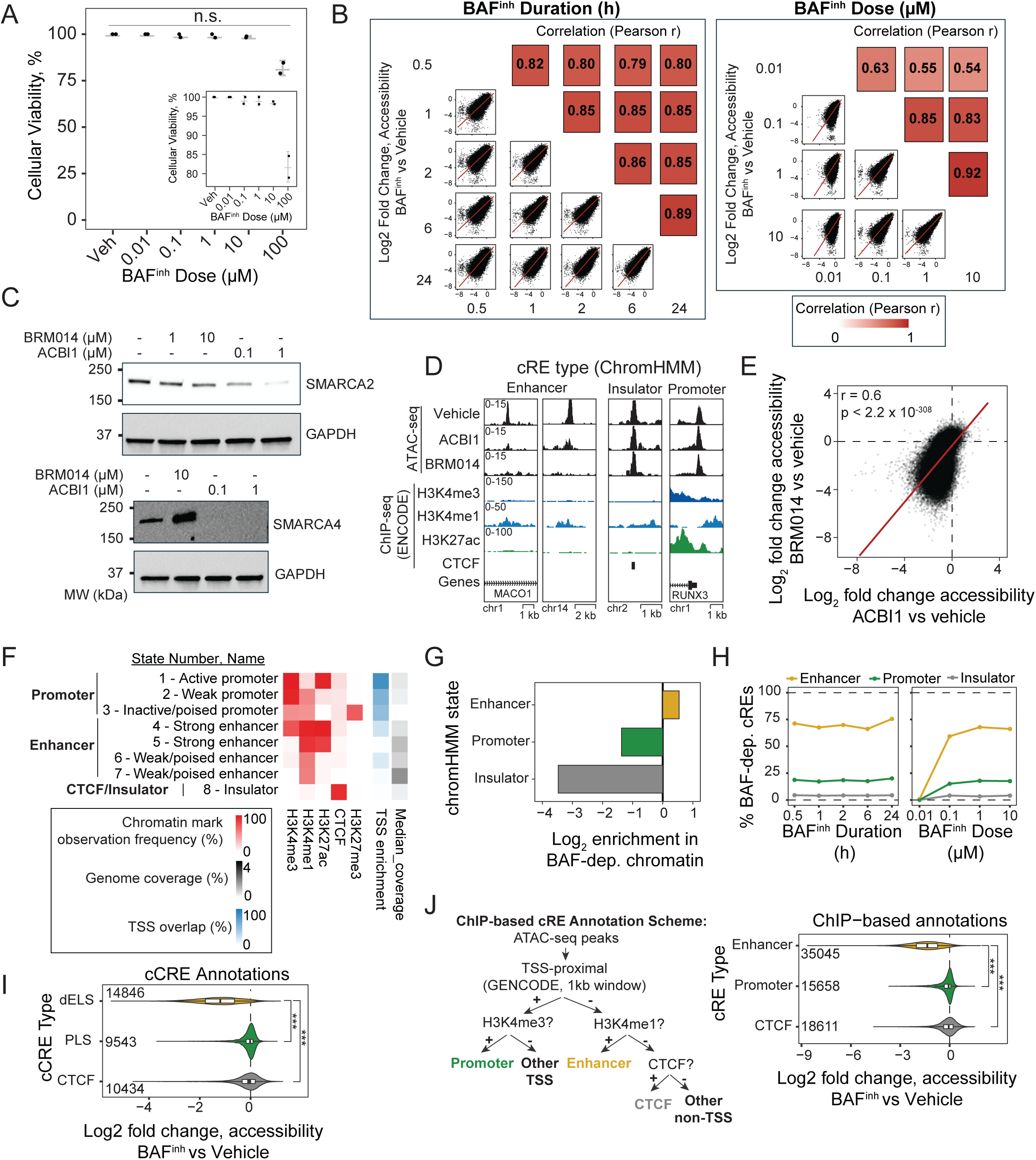
(Related to Figure 1): Pharmacological disruption of BAF activity results in widespread accessibility changes, with outsize effects at enhancers. (A) BRM014 treatment for 24 h does not significantly affect cell viability. The shows viability data with a restricted y-axis range. P-values were calculated using Student’s t-tests. n.s., not significant (B) Accessibility changes (from ATAC-seq) due to BAF^inh^ are strongly correlated across timepoints and doses. (C) Treatment with ACBI1, a proteolysis-targeting chimera against SMARCA2/A4, leads to undetectable SMARCA4 and strongly reduced SMARCA2 after 24h of treatment. A representative Western blot is shown. (D) Representative genome browser examples demonstrating similar effects of BAF^inh^ and ACBI1 at enhancers (left; decreased accessibility) and promoters and insulators (right; maintained accessibility). (E) Accessibility changes induced by BAF^inh^ and ACBI1 are strongly correlated. P-value calculated using a two-sided Pearson correlation test. (F) Enrichment of histone modifications and TSS enrichment between relevant ChromHMM states, adapted from Ernst et al., 2011 [25]. ChromHMM states used to define promoters, enhancers, and insulators are indicated at left. (G) Enhancer-associated ChromHMM states are enriched for BAF-dependent chromatin accessibility while promoter-and insulator-associated states are depleted. (H) Among cRE classes, enhancers show the highest proportion of significantly BAF-dependent elements across both BAF^inh^ dose and duration. (I) Using ENCODE candidate cis-regulatory element (cCRE) annotations [26], distal enhancer-like signature (dELS) cREs exhibit significantly greater accessibility loss upon BAF^inh^ compared to promoter-like signature (PLS) and CTCF-only elements. (J) Alternative cRE annotation scheme based on TSS proximity and ChIP–seq peak overlap (left). Enhancers defined using this approach similarly show greater loss of accessibility upon BAF^inh^ relative to other cRE classes. ***, p < 2.2 x 10^-308^, Wilcoxon rank-sum test.

**Supplemental Figure 2.**
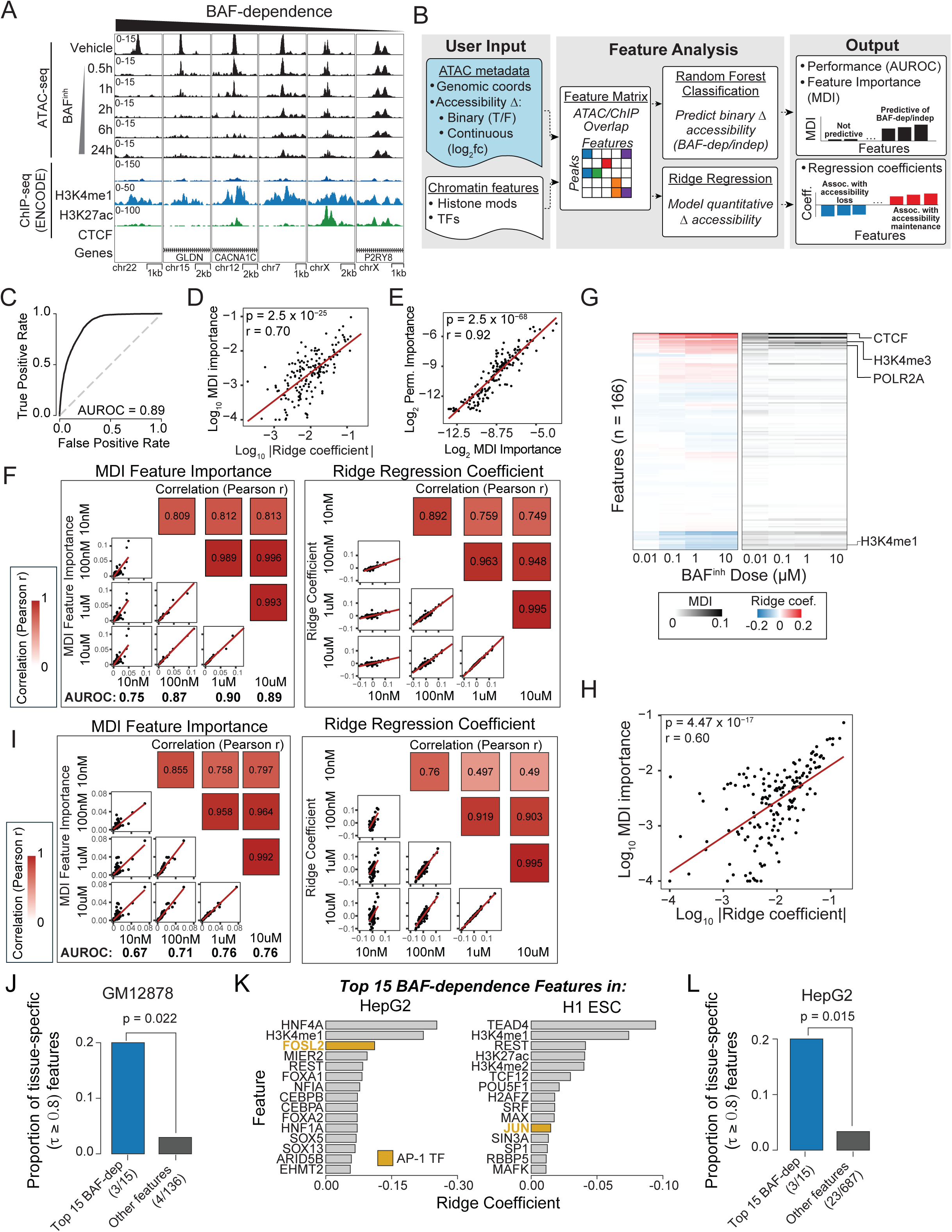
(Related to Figure 2): Robust performance and consistent feature associations in machine learning models predicting BAF-dependent accessibility. (A) Genome browser examples demonstrating heterogeneity in enhancer responses to BAF^inh^. (B) Overview of the machine learning approach. ENCODE TF and histone ChIP-seq experiments are filtered by experiment quality (FRiP score – see **Methods**) and intersected with consensus ATAC-seq peaks to construct a feature matrix, which is then used to model binary BAF dependence (random forest classification) and continuous accessibility changes (ridge regression). (C) Discriminative performance of the random forest model when applied to all cis-regulatory elements (cREs), assessed by AUROC. (D) Feature importance metrics derived from random forest (mean decrease in impurity) and ridge regression (regression coefficients) are highly correlated, demonstrating agreement between approaches. (E) MDI and permutation-based feature importances are strongly correlated. P-value calculated using two-sided Pearson correlation test. (F) Feature importance metrics from all-cRE models trained using different BAF^inh^ doses are highly correlated, indicating consistency of feature associations. (G) All-cRE models identify expected features associated with BAF-dependence (e.g., H3K4me1) and -independence (e.g., H3K4me3, Pol2, CTCF). (H) MDI feature importance and ridge regression coefficients calculated from an enhancer-only feature matrix are significantly correlated. (I) Feature importance metrics from enhancer-only models using different BAF^inh^ doses are highly concordant. (J) Features associated with BAF dependence are significantly enriched for tissue-restricted gene expression (τ > 0.8) compared to other features. P-value calculated using Fisher’s exact test; numbers of tissue-restricted and non-restricted features for each category are indicated (K) AP-1 TFs are represented among the top BAF-dependence features in both HepG2 and H1 embryonic stem cells. (L) In HepG2 cells, BAF-dependence features are significantly enriched for tissue-restricted expression. P-value calculated using Fisher’s exact test; numbers of tissue-restricted and non-restricted features for each category are indicated.

**Supplemental Figure 3.**
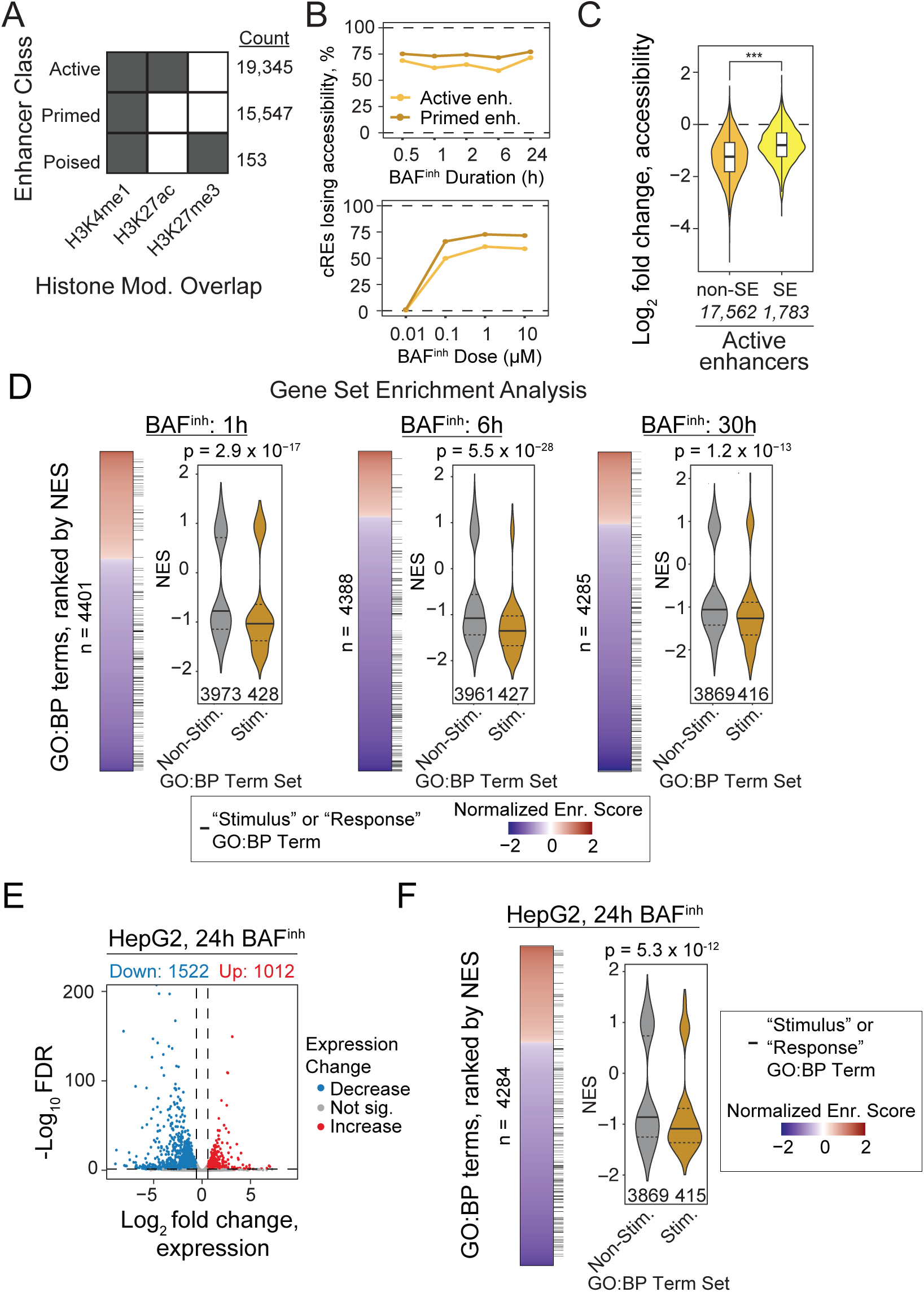
(Related to Figure 3): BAF^inh^ preferentially disrupts primed enhancers and stimulus-responsive transcriptional pathways. (A) Enhancers were classified as active, primed, or poised based on overlaps with H3K27ac and H3K27me3 ChIP-seq peaks. (B) Primed enhancers show a higher proportion of significantly BAF-dependent elements compared to active enhancers across both BAF^inh^ duration (top) and dose (bottom). (C) Active enhancers within super-enhancers (SEs) show reduced BAF-dependence compared with non-SE active enhancers. (D) Genes associated with stimulus–response pathways are downregulated following BAF^inh^ in GM12878 cells. Gene Ontology Biological Process (GO:BP) terms are ranked by Normalized Enrichment Scores (NES) from GSEA performed for 1h, 6h, and 30h BAF^inh^, displayed as heatmaps, and categorized based on whether term names contained “stimulus” or “response” (black bars). Violin plots show distributions of NES scores for stimulus-and non-stimulus term sets, where solid lines represent median NES and dashed lines represent 25^th^ and 75^th^ percentiles. P-values were calculated using one-sided Wilcoxon rank-sum tests; total numbers of GO:BP terms analyzed for each timepoint as well as sizes of stimulus and non-stimulus term sets are indicated. (E) Transcriptional changes upon BAF^inh^ (10µM, 24h) in HepG2 cells. (F) As in GM12878, stimulus-associated pathways are downregulated upon BAF^inh^ in HepG2. P-value calculated as in (D).

**Supplemental Figure 4.**
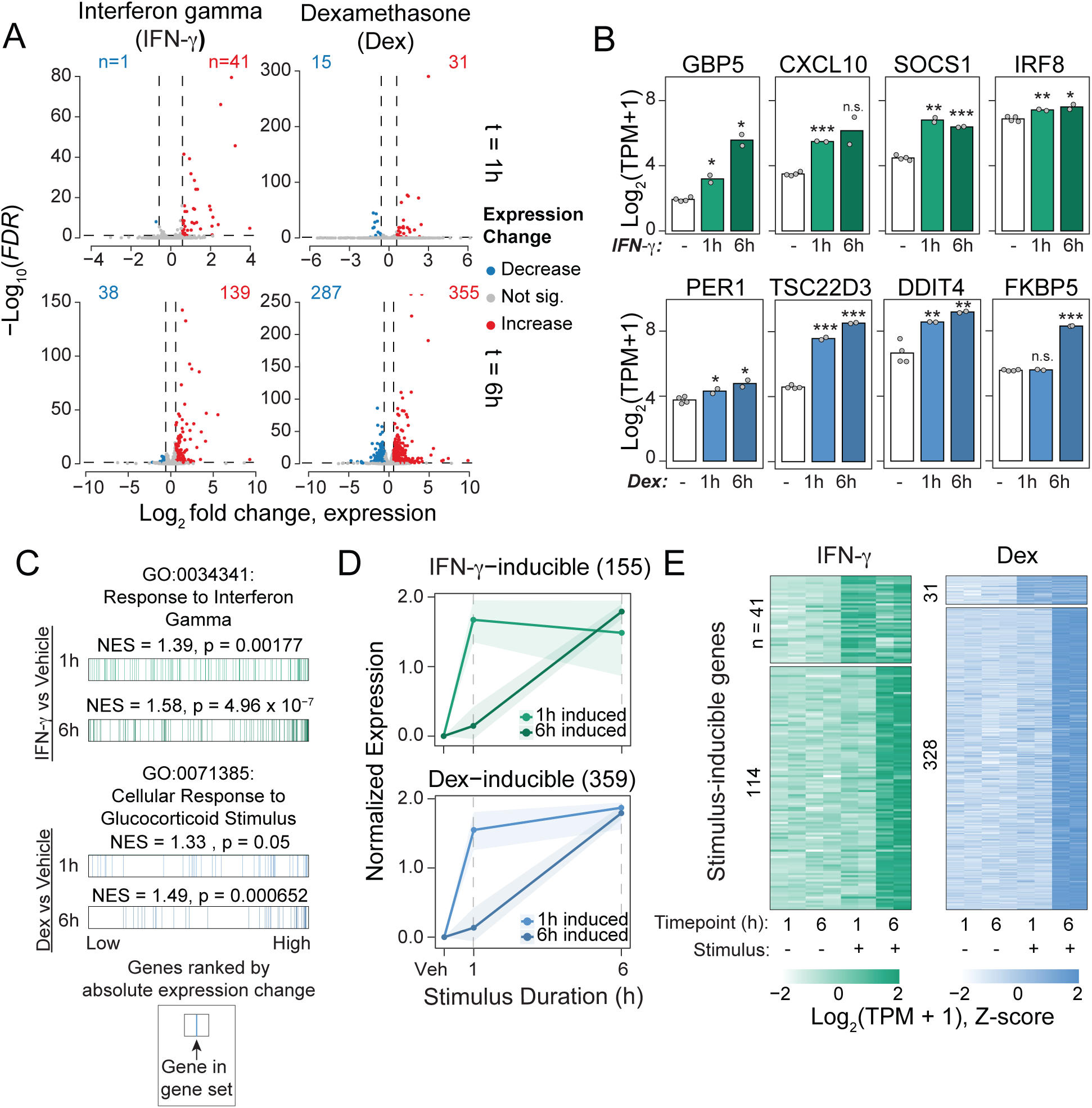
(Related to Figure 4): Time-dependent transcriptional responses to IFN-γ and dexamethasone in GM12878 cells. (A) Volcano plots showing the magnitude and direction of transcriptional responses to IFN-γ and dexamethasone (Dex) at 1 h and 6 h. Numbers of genes with significantly increased (FDR < 0.05, log2FC > log2(1.5)) or decreased (FDR < 0.05, log2FC < -log2(1.5)) expression are indicated. (B) Representative canonical IFN-γ– and Dex-induced genes, illustrating stimulus-specific transcriptional induction. P values were calculated using a Student’s *t*-test (*, p < 0.05; **, p < 0.01, ***, p < 0.001). (C) Specificity of transcriptional responses to IFN-γ and Dex demonstrated by gene set enrichment analysis (GSEA) on IFN-γ response and glucocorticoid response GO:BP terms, respectively. (D) Median expression trajectories of 1h- and 6h-induced genes following IFN-γ or Dex treatment. Expression is shown as Z-scored log2(TPM + 1), normalized to vehicle. Shaded regions denote the interquartile range. (E) Heatmap showing gene-level expression changes for early- and late-induced genes in response to IFN-γ and Dex.

**Supplemental Figure 5.**
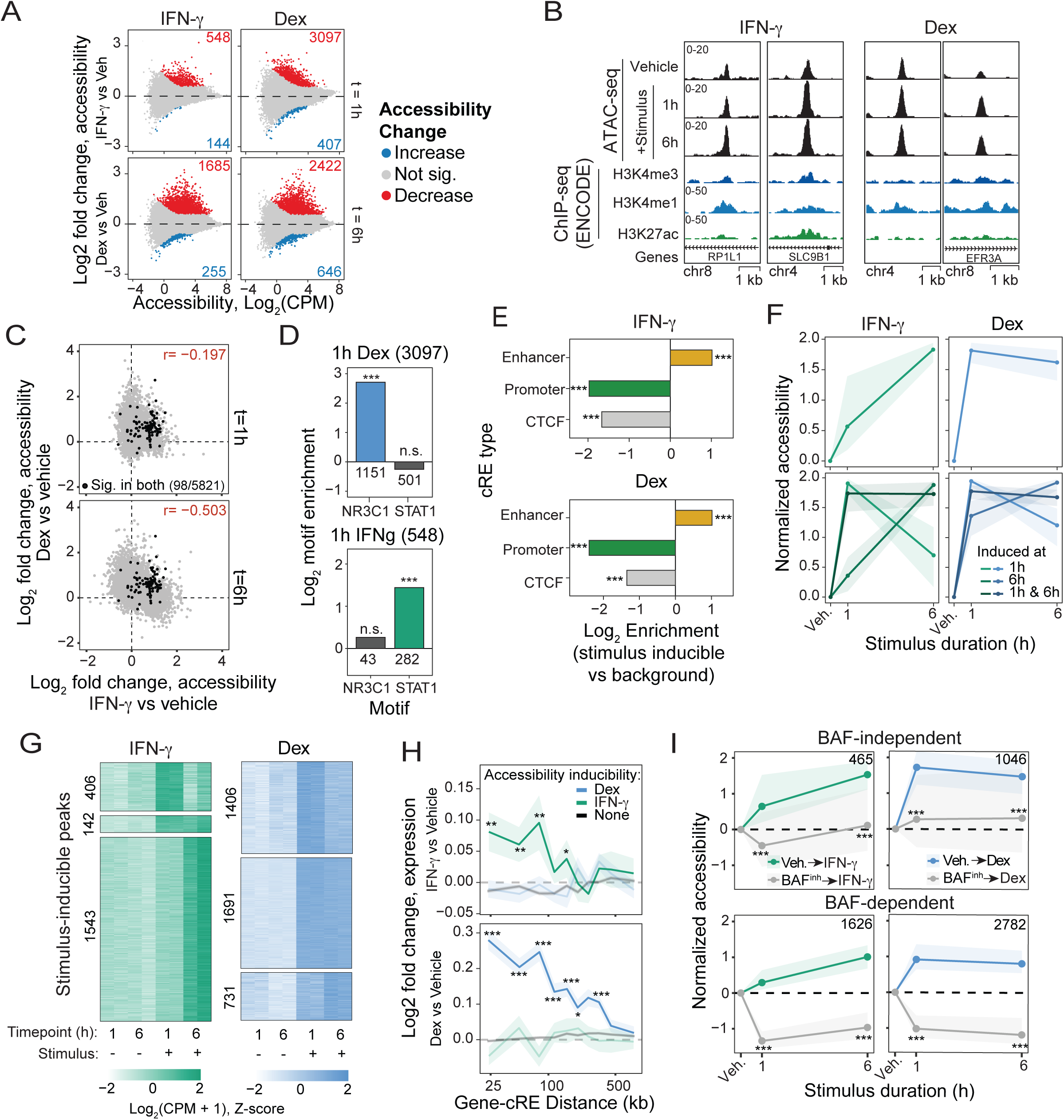
(Related to Figure 5): Chromatin accessibility changes in response to IFN-γ and dexamethasone in GM12878 cells. (A) MA plots showing chromatin accessibility changes in response to IFN-γ or Dex at 1 h and 6 h. (B) Representative genome browser examples of enhancers that gain accessibility in response to IFN-γ (left) or Dex (right). (C) Chromatin accessibility gains are stimulus specific. All cREs showing significant induction in either stimulus or time point are plotted by log2 fold change at 1 h (top) or 6 h (bottom); cREs significantly induced by both stimuli are indicated. (D Enriched transcription factor motifs at 1h stimulus-inducible cREs correspond to the applied stimulus. Numbers of inducible loci containing each motif are indicated. P values were calculated using Fisher’s exact tests (E) Stimulus-inducible cREs are enriched for enhancer annotations relative to background regions. P-values were calculated using Fisher’s exact tests (F) Median accessibility gains at all inducible regions (top) and separated by induction timepoint (1h-only, both timepoints, 6h-only). Median normalized accessibility (Z-scored log2(CPM+1), normalized to vehicle) is shown, with the shaded region representing the interquartile range. (G) For each IFN-γ (left) or Dex (right) inducible cRE, normalized accessibility levels are shown, separated by induction timepoint. cREs are grouped based on the timepoint(s) at which significant stimulus-responsive accessibility gains are observed: 1h-only (top), both timepoints (middle) and 6h-only (bottom). (H) Genes linked by promoter capture Hi-C (PCHi-C) to stimulus-inducible cREs exhibit stimulus-specific transcriptional responses. Genes linked to IFN-γ-inducible, Dex-inducible, or non-inducible cREs were binned by contact distance, and mean expression fold change in response to IFN-γ (top) or Dex (bottom) was calculated for each bin. Values are plotted as mean fold change (line) and standard error of the mean (shaded region). P values were calculated using Wilcoxon rank-sum tests (IFN-γ vs Dex) for each decile and adjusted for multiple testing using the Benjamini-Hochberg method). (I) After stratifying stimulus-inducible cREs by basal BAF dependence, both BAF-dependent and BAF-independent categories show reduced accessibility induction following BAF^inh^ pretreatment. Normalized accessibility and IQR is shown as in (G). Numbers of cREs in each category for each stimulus are indicated. P-values were calculated from Wilcoxon rank-sum tests. *, p < 0.05; **, p < 0.01; ***, p < 0.001; n.s., not significant.

